# PTEN Regulates Myofibroblast Activation in Valvular Interstitial Cells based on Subcellular Localization

**DOI:** 10.1101/2024.06.30.601424

**Authors:** Dilara Batan, Georgios Tseropoulos, Bruce E. Kirkpatrick, Kaustav Bera, Alex Khang, Mary Weiser-Evans, Kristi S. Anseth

## Abstract

Aortic valve stenosis (AVS) is characterized by altered mechanics of the valve leaflets, which disrupts blood flow through the aorta and can cause left ventricle hypotrophy. These changes in the valve tissue result in activation of resident valvular interstitial cells (VICs) into myofibroblasts, which have increased levels of αSMA in their stress fibers. The persistence of VIC myofibroblast activation is a hallmark of AVS. In recent years, the tumor suppressor gene phosphatase and tensin homolog (PTEN) has emerged as an important player in the regulation of fibrosis in various tissues (e.g., lung, skin), which motivated us to investigate PTEN as a potential protective factor against matrix-induced myofibroblast activation in VICs. In aortic valve samples from humans, we found high levels of PTEN in healthy tissue and low levels of PTEN in diseased tissue. Then, using pharmacological inducers to treat VIC cultures, we observed PTEN overexpression prevented stiffness-induced myofibroblast activation, whereas genetic and pharmacological inhibition of PTEN further activated myofibroblasts. We also observed increased nuclear PTEN localization in VICs cultured on stiff matrices, and nuclear PTEN also correlated with smaller nuclei, altered expression of histones and a quiescent fibroblast phenotype. Together, these results suggest that PTEN not only suppresses VIC activation, but functions to promote quiescence, and could serve as a potential pharmacological target for the treatment of AVS.

## Introduction

Aortic Valve Stenosis (AVS) is a progressive cardiovascular disease that directly affects the aortic valve leaflet matrix composition and mechanical properties, leading to disrupted blood flow through the aorta^1^ and eventual complications associated with left ventricular pressure-overload.^2^ When aortic valve leaflets thicken and stiffen as AVS progresses, significant deficiencies in cardiac output result, which can lead to heart failure and death. AVS affects around 2% of the population older than 65 with higher occurrences in males compared to females.^3^ Symptomatic AVS patients suffer high mortality rates if the condition is left untreated.^4^ Currently, there are no known effective pharmacological treatments for AVS, and surgical or transcatheter valve replacement remains the standard of care for patients with severe disease. However, many patients are ineligible for surgery, as valve replacement has many risks including valve leaks, bleeding, and vessel damage.^1,5^ As a result, a deeper molecular and mechanistic understanding of AVS is necessary if one hopes to identify alternative pharmaceutical treatment strategies.

Valvular interstitial cells (VICs) are the main population of cells residing in aortic valve leaflets, and these fibroblast cells are responsible for valve maintenance and repair following injury. Because of their key role in valve homeostasis, VICs are highly sensitive to physicochemical changes in the matrix microenvironment.^6–8^ Specifically, VICs transition from quiescent fibroblasts to activated myofibroblasts, characterized by formation of alpha-smooth muscle actin (αSMA) stress fibers, increased proliferation, and elevated extracellular matrix (ECM) deposition.^9^ During fibrosis, the myofibroblast phenotype persists over time, leading to a pathogenic accumulation of ECM, stiffening of the local microenvironment, and creating a positive feedback loop that can accelerate disease progression.^10^ Investigating mechanotransduction pathways capable of reversing this process and restoring the quiescent fibroblast phenotype continues to be a promising avenue for potential treatments.^11–15^

Hydrogels provide highly tunable microenvironments that can be adjusted to recapitulate the stiffness of the valve matrix at different stages of fibrosis progression and enable researchers to study VIC mechanosensing in a controlled manner.^16^ In this work, the crosslinking density of poly(ethylene glycol) (PEG) hydrogels was tailored to control the matrix mechanical properties,^17^ while the fibronectin-derived adhesive peptide, RDGS, was used to promote VIC-matrix interactions.^18^ When cultured on soft PEG hydrogels (∼1-5 kPa), VICs maintained a quiescent fibroblast phenotype, but transitioned to an activated myofibroblast phenotype on stiffer PEG matrices (E’∼13-30kPa).^15,19–22^ Thus, these biomaterial systems provided a reliable *in vitro* platform for VIC cultures in healthy and disease scenarios.

As valvular tissue stiffens during AVS progression, signaling through mechanosensitive pathways can reveal essential signaling hubs governing cell phenotype and potentially serve as druggable targets. Importantly, elevated levels of phospho-AKT (p-AKT), which indicates increased activation of phosphoinositide 3-kinase (PI3K)/AKT pathway, have been observed in the fibroblast to myofibroblast transition (FMT).^23^ In this regard, the pathway has been presented as a master regulator of idiopathic pulmonary fibrosis.^23^ Similarly, we have previously demonstrated that PI3K/AKT signaling is increased on stiff substrates and is necessary for substrate stiffness induced myofibroblast activation in VICs.^11,24^ These findings suggest that targeting the PI3K/AKT pathway may be a promising approach for treating fibrotic diseases.

Phosphatase and tensin homolog (PTEN), which is a negative regulator of the PI3K/AKT pathway, has been proposed as a potential target for therapeutic intervention in fibrotic disease.^25–27^ For example, Lu *et al.* demonstrated that overexpression of PTEN in the mouse model superPTEN protects against angiotensin II induced vascular fibrosis.^26^ Further, the same group later observed that small molecule drugs, such a 5-azacytidine and itraconazole, increase PTEN levels and are able to ameliorate effects of increased signaling through the PI3K/AKT pathway.^26,28^ Complementary to the overexpression studies, Parapuram *et al*. demonstrated that loss of PTEN leads to overproduction of collagen and is enough to induce fibrosis in lung and skin tissues.^27,29^ Similar studies were conducted in mammary gland tissue by Jones *et al.* and further confirmed that PTEN regulates extracellular matrix deposition and collagen organization, steps that are essential in fibrosis development.^30^ Collectively, these findings suggest that targeting the PI3K/AKT through PTEN may hold promise for developing novel therapies for fibrotic diseases. However, no previous work has been conducted specifically investigating the effects of PTEN in the context of AVS. Motivated by this, we aimed to explore the role of PTEN in VIC myofibroblast activation and its potential as a therapeutic target for aortic valve disease.

Results herein reveal a major role of PTEN in regulating VIC fibroblast and myofibroblast phenotypes. Initially, we observe a loss of PTEN protein in diseased human valve leaflet sections compared to healthy ones. To test the specific role of mechanotransduction on PTEN expression and VIC phenotype, we tailored the modulus/stiffness of PEG hydrogels to direct VICs towards either a fibroblast (E’ ∼3 kPa) or a myofibroblast (E’ ∼13.4 kPa) majority population. Next, we demonstrate that pharmacologically increased PTEN levels promote a more quiescent phenotype for VICs cultured on stiff hydrogels, while chemical and genetic inhibition of PTEN increased myofibroblast activation on soft hydrogels. Interestingly, the minority fibroblast population on stiff matrices had higher nuclear localization of PTEN compared to the majority myofibroblast population. In addition, the population of cells with high PTEN nuclear localization correlated with smaller nuclei and VIC quiescence. The regulation of VIC activation by PTEN was further verified by shRNA knockdown, as well as overexpression (nuclear localization and exclusion signal tagged) experiments *in vitro*. Our results led us to hypothesize that nuclear PTEN may play an additional role, beyond cytoplasmic regulation of PI3K/AKT, maintaining VIC quiescence by regulating histone expression on stiff matrices. Overall, in this study we show that PTEN is a major regulator of VIC fibroblast to myofibroblast transition and can serve as a future target for pharmaceutical interventions.

## Materials and Methods

### Hydrogel precursor synthesis

Poly(ethylene glycol) photodegradable diacrylate PEGdiPDA (MW approximately 3400 Da) was synthesized and characterized according to a previous method.^31^ >90% acrylate functionalization was verified by ^1^H NMR. The adhesive peptide, acrylamide diethylene glycol-diethylene glycol-glycine-arginine-glycine-aspartic acid-serine-glycine (Ac-OOGRGDSG), was synthesized via 9-fluorenylmethyloxycarbonyl (Fmoc)/1-[bis(dimethylamino)methylene]-1H-1,2,3-triazolo[4,5-b] pyridinium 3-oxid hexafluorophosphate (HATU) coupling strategy on Rink Amide methylbenzhydryl amine (MBHA) resin (0.500 mmol) using a Protein Technologies Tribute automated solid-phase peptide synthesis machine and characterized using electrospray ionization (ESI) mass spectroscopy.^31^

### Formation of hydrogels

The preparation of photo-softening PEGdiPDA hydrogels was adapted from previously described protocols.^31^ PEGdiPDA was co-polymerized with poly(ethylene glycol) monoacrylate (PEGA; Mn∼400 Da; Monomer-Polymer and Dajac Laboratories) and acryl-OOGRGDSG in PBS via redox-initiated free radical polymerization. Gel solutions were prepared with 7.0 wt% PEGdiPDA, 6.8 wt% PEGA, 5 mM acryl-OOGRGDSG, 0.2 M ammonium persulfate (APS) and 0.1 M tetramethylethylenediamine (TEMED). Gels were formed on acrylated cover glass after the addition of APS/TEMED, with diameters of 12, 15, or 25 mm and a thickness of 100 µm. Gels were allowed to cure for 6 min and washed and stored in PBS before cell seeding. Soft hydrogels were prepared by irradiating the initial hydrogels with UV light (λ = 365 nm; I_0_ = 10 mW/cm^2^) for 6 minutes.

### Swollen rheological characterization

Optically thin PEGdiPDA hydrogels (thickness = 200 μm) were made between two sigmacoted glass slides separated with a spacer using the formulations above and allowed to swell in PBS overnight. A rheometer (TA Instruments DH-R3) with a parallel plate accessory was used to measure the shear modulus of the PEGdiPDA hydrogel before and after photo-degradation. The equilibrium storage modulus (G′) was measured using a dynamic time sweep (γ = 1%; ω = 1 rad/s), as determined to be in the linear regime for this material. For *in situ* degradation, a quartz plate was used, and the rheometer was equipped with a 365 nm light source (λ = 365 nm; I_0_ = 10 mW/cm^2^; Omnicure 1000, LumenDynamics). After network formation, the hydrogel was exposed to 365 nm light for 6 minutes, and the change in G’ was monitored using the same dynamic time sweep parameters. The same dynamic time sweep measurements were used to obtain final G’ as a function of light exposure time. G’ was measured and converted to Young’s modulus (E) using a Poisson’s ratio of 0.5.^32^

### Valvular Interstitial Cell (VIC) isolation and culture

Primary VICs were isolated from aortic valve leaflets from fresh male porcine hearts (Hormel) as previously described.^14,20,21,24^. Briefly, aortic valve leaflets were excised from the hearts and rinsed in Earle’s Balanced Salt Solution (EBSS, Life Technologies) supplemented with 50 U/mL penicillin and 50 µg/mL streptomycin (Thermo Fisher), and 1 µg/mL amphotericin B (Thermo Fisher). Leaflets were incubated in a 250 U/mL type II collagenase (Worthington) solution in EBSS for 30 min under 5% CO2 at 37°C with gentle shaking. Valvular endothelial cells were removed by vortexing leaflets at maximum speed for 30 seconds. Leaflets were transferred to fresh collagenase solution and incubated for 60 min under 5% CO2 at 37°C with gentle shaking. Leaflets were vortexed again for 2 minutes to dislodge valvular interstitial cells (VICs), and the entire solution was passed through a 100 µm cell strainer. The strained cell solution was centrifuged at 0.2g x 20 minutes. Pelleted cells were resuspended in VIC expansion media compromising of M199 media (Life Technologies) 15% fetal bovine serum (FBS, Life Technologies), 50 U/mL penicillin, 50 µg/mL streptomycin (Thermo Fisher), and 1µg/mL amphotericin B (Thermo Fisher). Cells were cultured in T-75 flasks for expansion until 80% confluency was obtained. Cells were harvested using trypsin (Life Technologies) to be frozen in 50% fetal bovine serum (FBS) (Life Technologies), 45% M199 media (Life Technologies), and 5% DMSO (Sigma-Aldrich).

For experimental use, cells were thawed and cultured in T-75 flasks in 15% FBS supplemented M199 media until 80% confluent. Cells were counted after harvesting using an automated hemocytometer (Countess). A seeding density of 15,000 cells/cm^2^ was used for VICs on hydrogels, with growth media consisting of M199 media supplemented with 1% FBS, penicillin-streptomycin, and fungizone. 12 mm hydrogels were used for immunostaining experiments; 25 mm hydrogels were used for real-time quantitative polymerase chain reaction (RT-qPCR) to increase the measurable RNA content.

### Hydrogel Immunostaining

VICs were fixed in 4% paraformaldehyde (PFA, Electron Microscopy Sciences) in PBS for 20 minutes and rinsed twice with PBS. Cells were permeabilized with 0.1% TritonX-100 (Fisher Scientific) in PBS for 20 minutes followed by 1 hour of blocking in 5% bovine serum albumin (BSA, Sigma-Aldrich). Primary antibodies were diluted in 5% BSA and incubated at 4°C overnight at given concentrations: α-smooth muscle actin (αSMA, mouse, Cat. No. ab7817, Abcam) at 1:300 dilution, PTEN (rabbit, Cat. No. D8H1X, Cell Signaling) at 1:300 dilution. Primary antibody was washed off the cells twice with PBST (0.5% Tween-20 in PBS), and once in PBS. Samples were incubated in secondary antibody solution of 1:200 dilution goat anti-mouse Alexa 488 (Life Technologies), 1:500 dilution of goat anti-rabbit Alexa 648 (Life Technologies), 1:5000 dilution of HCS Cell Mask (Life Technologies), and 1:1000 dilution of DAPI (Life Technologies) for 1 hour before rinsing twice with PBST (0.005% Tween-20 in PBS) and once in PBS. For imaging, hydrogels were inverted onto a glass-bottom 24-well plate (Cellvis) and visualized using an Operetta high-content confocal microscope using a 20x air objective (Perkin Elmer). Immunofluorescence intensity in immunostained cells was quantified using Harmony software (Perkin Elmer). PTEN nuclear localization was quantified by dividing PTEN average nuclear signal by PTEN average cytoplasmic signal.

### Isolation of mRNA and Real-Time Quantitative Polymerase Chain Reaction (RT-qPCR)

RNA was isolated according to manufacturer’s instructions using an RNeasy Micro Kit (Qiagen). RNA concentration was quantified using ND-1000 NanoDrop Spectrophotometer (Thermo Fischer). cDNA was produced using iScript Reverse Transcription Supermix kit (Bio-Rad) on a Mastercycler (Eppendorf). Relative mRNA levels were measured with SYBR Green reagents on an iCycler (Bio-Rad). Results were normalized to the housekeeping gene RPL30 (Ribosomal protein L30). Custom primers are presented in **Table S1**.

### PTEN lentiviral knockdown

Two shRNA sequences were purchased from Millipore Sigma and cloned into the pLKO.1 lentiviral vector. Virus was produced utilizing second generation packaging plasmids PSPAX2 and pnd2.g as described previously.^33^ Briefly, HEK 293T cells were transfected with the packaging plasmids and shRNA for PTEN and two viral batches were collected after 24 and 48 hours. P1 VICs were transduced with the viral constructs for 3 hours, and VICs that incorporated the virus were selected with puromycin (1ug/ml) for 3 days. The genetically modified VICs were allowed to proliferate on TCPS until 80% confluence and subsequently passaged on soft or stiff hydrogels (E’ ∼3 kPa, E’ ∼13.4 kPa, respectively). The potency of shRNA PTEN knockdown was validated though immunohistochemistry for PTEN staining and qPCR testing for PTEN mRNA, and the most potent knockdown sequence was selected for experiments reported herein.

### Transfection with NES and NLS plasmid constructs

VICs were seeded on stiff hydrogels at a density of 6000 cells / cm^2^ and incubated overnight. Transfection media was prepared the next day using lipofectamine 3000 at a dilution of 0.5ul in 50ul of Opti-MEM reduced serum media per well and vortexed. The plasmids (Addgene PTEN NES Plasmid# 10932, Addgene PTEN NLS Plasmid# 10933) were amplified using the Qiagen^TN^ mini prep kit and diluted in a concentration of 1ug/ul. The control condition consisted of lipofectamine and Opti-MEM only. The growth media was removed from wells and transfection media was added. The transfection plate was incubated for 1 hour and then washed three times with Opti-MEM. Subsequently, growth media was added and the plate was incubated for 3 days, after which the plate was fixed and prepared for immunohistochemical characterization.

### Heart valve tissue histology

Samples of aortic healthy and diseased (AVS) valve leaflets were acquired from deceased or heart transplant patients at the University of Colorado Anschutz Medical Campus **(Table S2)**. Tissues were fixed in 4% paraformaldehyde for 10 minutes and embedded in Tissue Tek O.C.T. for subsequent cryosectioning. The sections were washed 3 times with PBS and permeabilized with 0.1% TritonX for 1 hour and blocked with 5% goat serum, 5% BSA, and 0.01% TritonX for 1 hour. The sections were treated with the following primary antibodies (αSMA, mouse, Cat. No. ab7817, Abcam at 1:300 dilution, PTEN rabbit, Cat. No. D8H1X, Cell Signaling at 1:300 dilution) sequentially over 2 days with 2 overnight incubations with the primary antibody in blocking buffer (dilution 1:200) and 1 hour incubation with secondary antibody (1:400). The tissues were imaged on an Nikon A1R confocal microscope and analyzed using ImageJ.

### Statistical Analysis

Data are presented as mean ± standard deviation with a minimum of three biological and four technical replicates for all studies unless otherwise stated. Outliers were removed using Robust Outlier (ROUT) method (set at 1%). Significance was defined as *P<0.05, **P<0.01, ***P<0.005, ****P<0.001 using a t-test for comparison of two samples and one-way analysis of variance (ANOVA) for comparison of three or more samples. Statistics were performed in GraphPad Prism.

## Results

### PTEN levels are lower in diseased human valve leaflets compared to healthy valves

As previous studies have demonstrated that loss of PTEN is involved in fibrosis development, we wanted to test whether PTEN levels were changed with AVS development. To do this, we immunostained human valve leaflet tissue sections from healthy and diseased patients for DAPI, αSMA, and PTEN and performed histological analysis targeted towards the levels of αSMA and PTEN. In healthy tissue, we observed high levels of PTEN and low levels of αSMA (**Figure 1 A-C**). In diseased tissue, significantly higher levels of αSMA and minimal levels of PTEN were observed, indicating an inverse correlation, and implicating a protective role for PTEN against fibrosis development in valve leaflets (**Figure 1 A**). These results motivated us to investigate the role of PTEN in VIC activation *in vitro,* building on previous work that utilizes soft and stiff hydrogels to recapitulate VIC myofibroblast differentiation and differential expression of αSMA, dependent on substrate stiffness.

**Figure 1:**
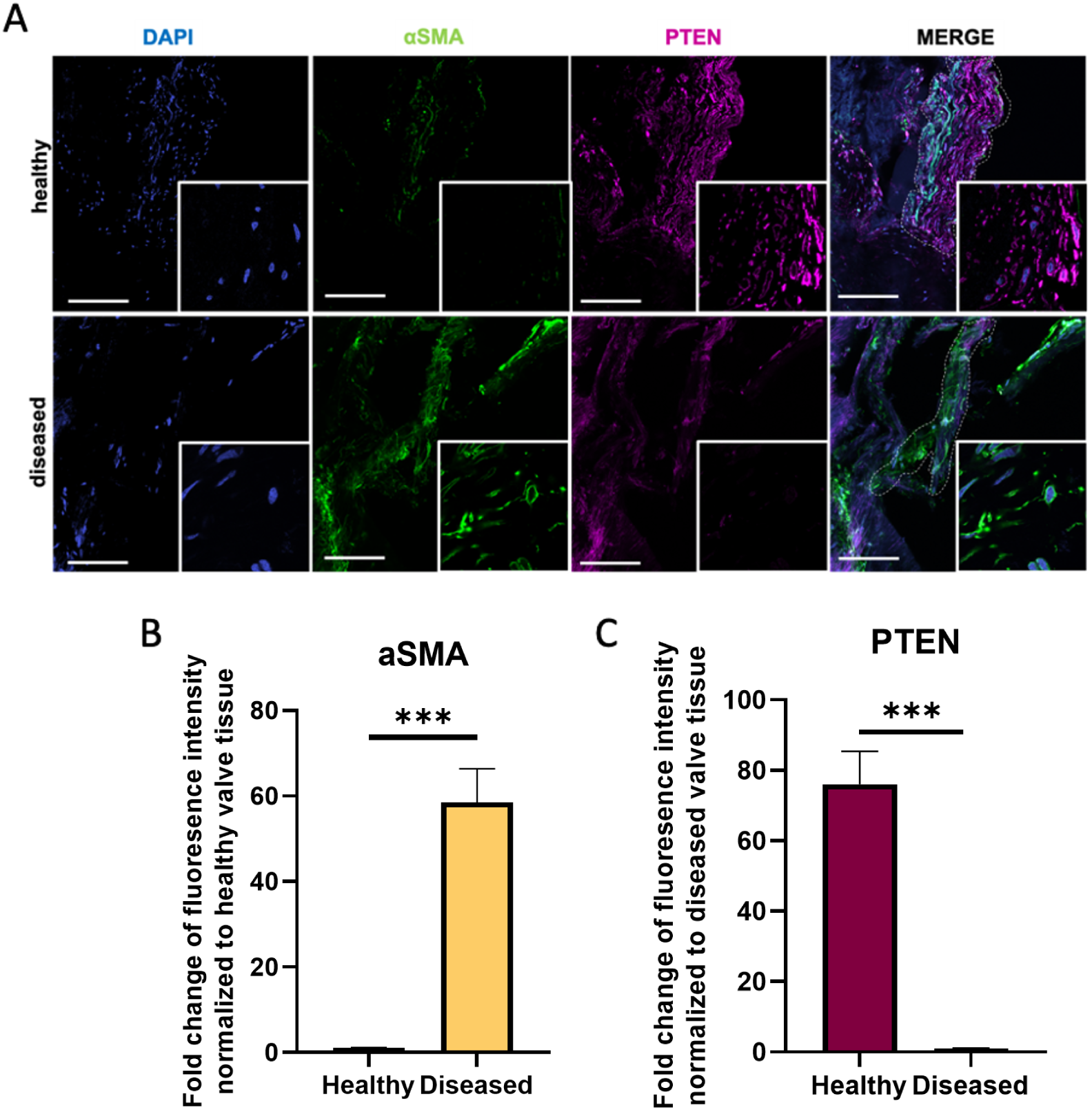
PTEN and *α*SMA levels inversely correlate in healthy and AVS human aortic valve tissue. **A.** Representative histological images of AVS and healthy human aortic valve tissue (male) stained for DAPI (blue), PTEN (magenta) and αSMA (green) levels in valvular interstitial cells. Scale bar = 50 µm **B.** Quantification of average αSMA intensity (normalized to DAPI per mm^2^) for healthy and diseased tissue within the marked area in A. (normalized to DAPI per mm^2^, n=5 fields of view per tissue, t-test, ***p<0.001). C. Quantification of PTEN intensity (normalized per mm^2^) for healthy and diseased tissue within the marked area. (n=5 fields of view per tissue, t-test, ***p<0.001)

### PTEN levels inversely correlate with αSMA levels in VIC hydrogel cultures

Motivated by the results from human valve samples, we wanted to test the effect of substrate stiffness on PTEN expression levels. To do this, we initially cultured VICs on soft and stiff hydrogels to identify conditions which promote a majority fibroblast phenotype on soft hydrogels and a majority myofibroblast phenotype on stiff hydrogels. We synthesized PEG-acrylate hydrogels supplemented with a fibronectin-derived peptide sequence, arginine-glycine-aspartic acid-serine (RGDS) to allow for cell adhesion and facilitation of cell-matrix interactions. We formulated soft (Young’s modulus (E’) = ∼2.8 kPa) and stiff hydrogels (Young’s modulus (E’) = ∼13.4 kPa) to match the modulus of healthy and diseased valve leaflet tissue to mimic the local microenvironmental VIC mechanosensing in AVS **(Supplementary Figure 1)**. Consistent with previous data, we observed a quiescent fibroblast VIC phenotype on soft hydrogels and an activated myofibroblast phenotype on stiff gels, as identified by the presence and intensity of αSMA **(Figure 2 A,B)**. In addition to the αSMA protein produced, the VIC morphology cultured on soft hydrogels was significantly smaller in cell and nuclear area compared to VICs cultured on stiff hydrogels **(Supplementary Figure 1 B-C)**. Next, to determine if PTEN plays a protective role against VIC myofibroblast activation, we examined protein levels in VICs cultured on the same soft and stiff hydrogels. We observed that VICs displayed significantly higher PTEN levels when cultured on soft hydrogels compared to stiff hydrogels, inversely correlating with αSMA levels and myofibroblast activation **(Figure 2 C,D)**.

**Figure 2:**
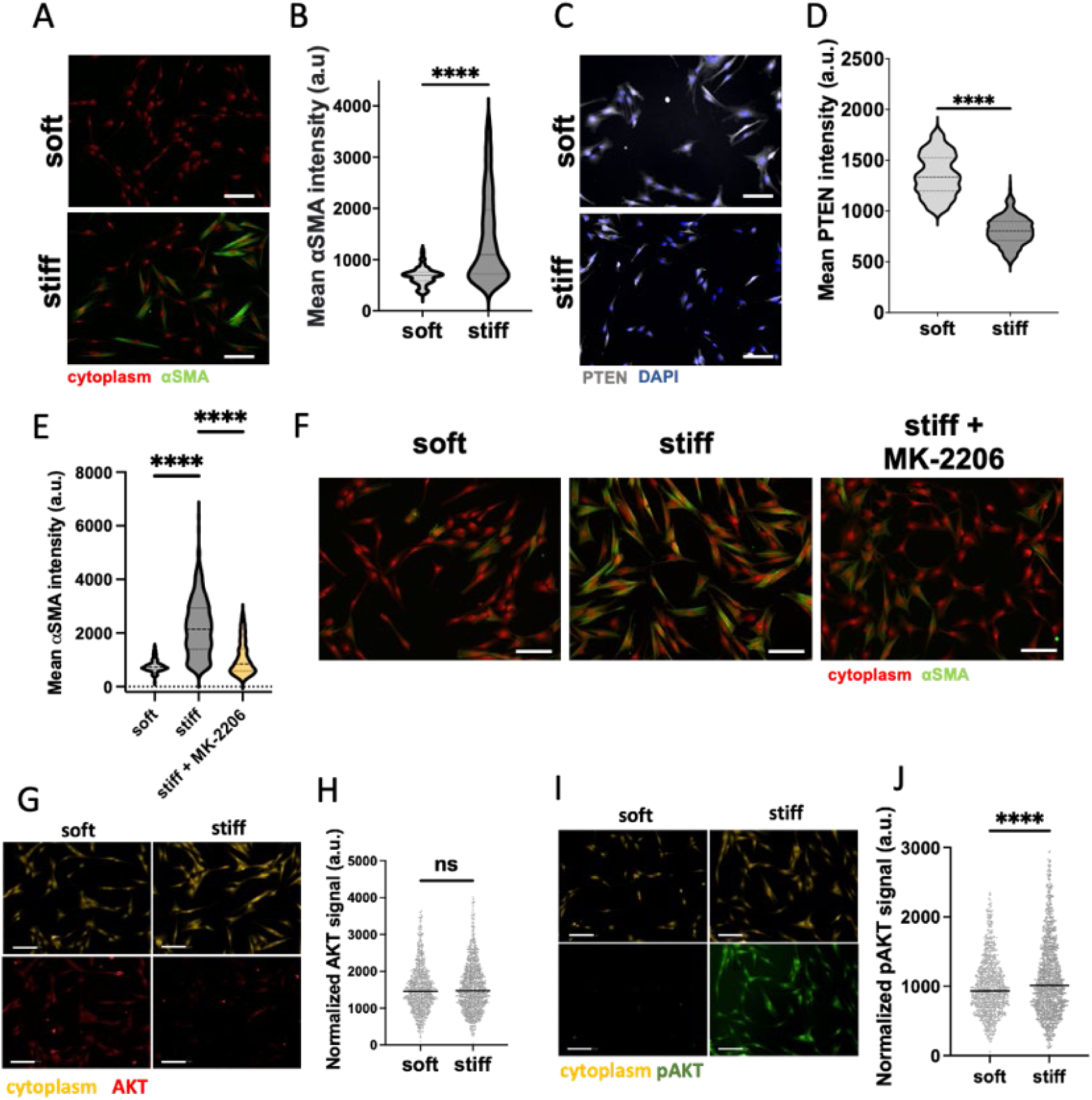
PTEN levels inversely correlate with myofibroblast activation in VICs. **A.** Representative images of VICs cultured on soft and stiff hydrogels for 3 days, αSMA (green) and cytoplasm (red). Scale bar = 100 µm **B.** Quantification of αSMA (green) intensity for A. (n>750 cells from 4 hydrogels, unpaired t test, ****p<0.001). **C.** Representative images of PTEN (gray) levels in VICs cultured on soft and stiff hydrogels for 3 days. **D.** Quantification of PTEN intensity for C. (n>440 cells from 3 hydrogels, unpaired t test, ****p<0.001). **E.** Quantification of αSMA (green) intensity for F. **F.** Representative images of VICs cultured on soft and stiff hydrogels for 3 days treated with DMSO vehicle or 5 µM AKT inhibitor MK-2206, αSMA (green) and cytoplasm(red). Scale bar = 100 µm (n>750 cells from 4 hydrogels, unpaired t test, ****p<0.001). **G.** Representative images of VICs cultured on soft and stiff hydrogels for 3 days, AKT (red) and cytoplasm (yellow). Scale bar = 100 µm **H.** Quantification of AKT (red) intensity for G. (n>920 cells from 3 hydrogels, unpaired t test). **I.** Representative images of VICs cultured on soft and stiff hydrogels for 3 days, pAKT (green) and cytoplasm (yellow). Scale bar = 100 µm **J.** Quantification of pAKT (green) intensity for I. (n>1000 cells from 3 hydrogels, unpaired t test, ****p<0.001).

Since PTEN is a known inhibitor of the PI3K/AKT pathway, we sought to verify its potential role in influencing stiffness-induced PI3K/AKT activity in VICs myofibroblast cultured on soft and stiff hydrogels. Specifically, we used a highly specific and widely used small molecule AKT inhibitor, MK-2206, and introduced it to activated VICs cultured on stiff hydrogels.^34–36^ We found that treatment with 5µM MK-2206 significantly reduced VIC αSMA expression when cultured on stiff hydrogels compared to untreated controls, and promoted a quiescent fibroblast population **(Figure 2 E,F, Supplementary Figure 2)**. Further, we observed similar levels of AKT in VICs cultured on soft and stiff hydrogels; however, pAKT levels were significantly higher in VICs cultured on stiff hydrogels (**Figure 2 G-J**). This observation supports the notion that PI3K/AKT is a key mechanoregulator of VIC myofibroblast activation in stiff microenvironments.^24^

### PTEN activators decrease VIC activation in stiff hydrogel microenvironments

Next, we wanted to investigate if increasing PTEN levels in VICs cultured on stiff hydrogels would play a protective role against myofibroblast activation. To test this, we used two small molecule PTEN activators that have been shown to increase cellular PTEN levels.^28,37^ Initially, we treated VICs cultured on TCPS with varying doses of the PTEN activators, 5-azacytidine or itraconazole, and compared these to the DMSO vehicle control. We tested PTEN mRNA levels using a concentration range (100 nm, 1 µM, and 10 µM) of 5-azacytidine and itraconazole and observed that PTEN levels were significantly increased with 10 µM treatment of both 5-azacytidine (**Figure 3A**) and itraconazole (**Supplementary Figure 3**).

**Figure 3:**
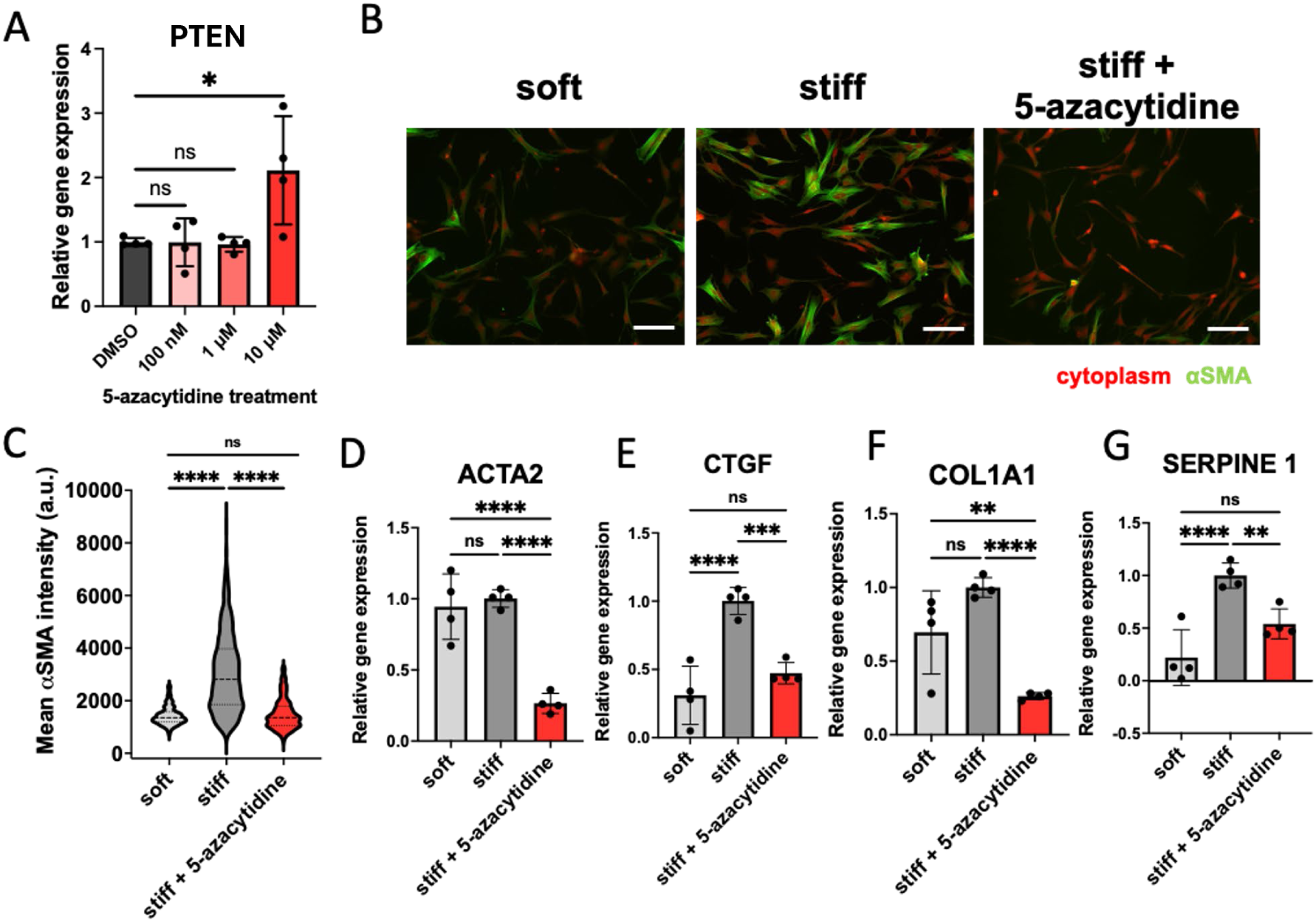
PTEN activators decrease myofibroblast activation in VICs. **A.** Relative mRNA expression levels of PTEN in VICs cultured on TCPS with treatment of vehicle or increasing concentration of the PTEN activator, 5-azacytidine, n=4 wells, means ± SD shown, one-way ANOVA test, *p < .05, **p < .01. **B.** Representative images of VICs cultured on soft and stiff hydrogels for 3 days in the presence of vehicle or PTEN activator 5-azacytidine, αSMA (green) and cytoplasm (red). Scale bar = 100 µm. **C.** Quantification of αSMA intensity for B (n>600 cells from 4 hydrogels, one-way ANOVA test, ****p<0.001). **D-G.** Relative mRNA expression levels of fibrotic genes in VICs cultured on soft or stiff hydrogels treated with vehicle or PTEN activator 5-azacytidine for 3 days, n=4 hydrogels, one-way ANOVA test, *p < .05, **p < .01, ***p < .001,****p<0.001.

After verifying that PTEN levels could be elevated using small molecule activators, we treated VICs cultured on stiff hydrogels with either vehicle or 5-azacytidine and itraconazole with previously determined concentrations **(Supplementary Figure 4 A-D)**. We observed that VICs treated with PTEN activators displayed significantly lower levels of αSMA compared to the vehicle treated stiff hydrogel culture controls at levels resembling the soft hydrogel negative controls **(Figure 3 B,C, Supplementary Figure 5 A,B)**. In addition to having lower levels of αSMA, VICs treated with PTEN activators also displayed a significantly smaller cell size and nuclear area when compared to DMSO treated controls **(Supplementary Figure 6 A,B)**.

To more broadly assess the role of PTEN in myofibroblast activation, VICs were cultured in the presence of PTEN activators or DMSO vehicle, and qPCR was performed to determine the relative expression of myofibroblast associated genes, namely alpha-Smooth Muscle Actin (ACTA2), Connective Tissue Growth Factor (CTGF), Serpin Peptidase Inhibitor clade E member 1 (SERPINE1), and Collagen type 1A1 (COL1A1). The expression of all four genes was significantly decreased in VICs cultured on stiff hydrogels treated with PTEN activators compared to the vehicle controls at levels resembling the soft hydrogel negative control conditions (**Figure 3 D-G**, **Supplementary Figure 5 C,D)**. These results suggest that PTEN influences multiple markers of VIC myofibroblasts and that the effects are observable after 72 hours of treatment.

### Chemical and genetic PTEN inhibition increases VIC myofibroblast activation in soft hydrogel microenvironments

To further validate the role of PTEN in myofibroblast activation, we next performed a series of studies inhibiting PTEN activity by introducing chemical and genetic perturbations. Initially we tested the effects of a potent PTEN phosphatase inhibitor bpV(HOpic) on VIC myofibroblast activation, as measured by αSMA expression.^38^ Quiescent VICs cultured on soft hydrogels treated with 100 nM of bpV(HOpic) displayed significantly higher levels of αSMA protein compared to cells treated with the DMSO control **(Figure 4 A,B)**. In addition to αSMA protein content, ACTA2 gene expression levels were also elevated with PTEN inhibitor treatment **(Figure 4 C)**. These results suggest that inhibition of PTEN phosphatase activity alone can promote myofibroblast activation in VICs. However, interestingly, we observed non-significant changes in mRNA levels of COL1A1 and SERPINE 1, suggesting more than just phosphatase activity of PTEN is important to induce these transcriptional changes (**Supplementary Figure 7**).

**Figure 4:**
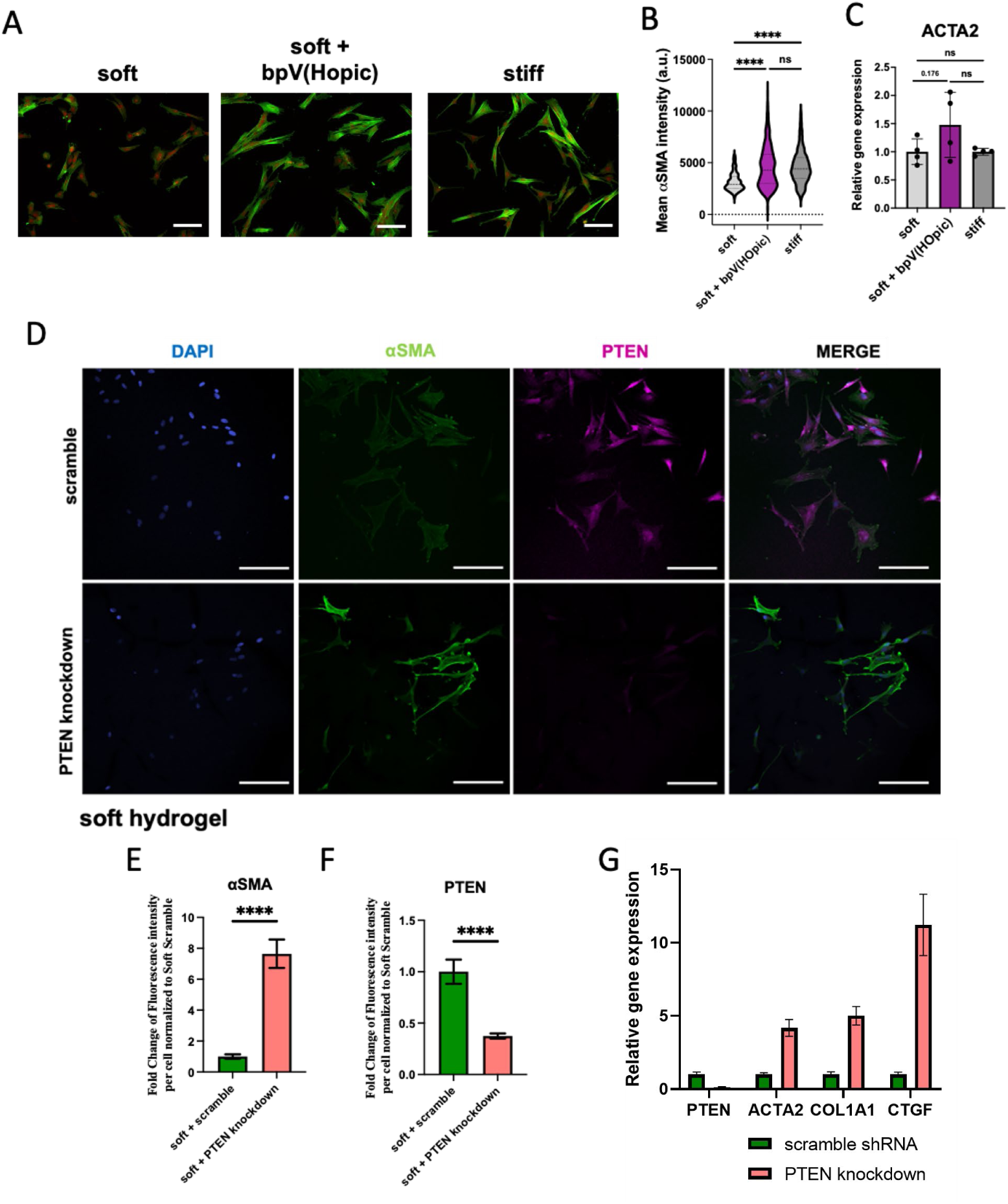
PTEN inhibitors increase myofibroblast activation in VICs. **A.** Representative images of VICs cultured on soft and stiff hydrogels for 3 days in the presence of DMSO vehicle or 100nM of PTEN inhibitor bpV(Hopic), αSMA (green) and cytoplasm(red). Scale bar = 100 µm. **B.** Quantification of αSMA intensity for A. (n>600 cells from 4 hydrogels, one-way ANOVA test, ****p<0.001). **C.** Relative mRNA expression levels of ACTA2 in VICs cultured on soft hydrogels with treatment of DMSO vehicle or 100nM of PTEN inhibitor bpV(Hopic), and stiff hydrogels, n=4 wells, means ± SD shown. **D.** Representative images of scrambled and PTEN knockdown VICs cultured on stiff hydrogels for 3 days, αSMA (green) and cytoplasm(red). Scale bar = 100 µm. **E.** Quantification of αSMA fluorescence intensity (n>30 cells, t-test, ****p<0.001) **F.** Quantification of PTEN fluorescence intensity (n>30 cells, t-test, ****p<0.001). **G.** Relative mRNA expression levels of various fibrotic genes in VICs treated with the scrambled or PTEN on TCPS, n=1 vector.

In addition to chemical PTEN inhibition, we utilized lentiviral shRNA knockdowns of PTEN in transduced early passage VICs. PTEN protein levels in the knockdown VICs were significantly reduced compared to the scrambled shRNA controls on both soft and stiff hydrogels. **(Figure 4 D,F, Supplementary Figure 8 A,C)**. We cultured scrambled or PTEN knockdown VICs on both soft and stiff hydrogels. On soft hydrogels, consistent with our previous data, we observed enhanced PTEN expression and low levels of VIC myofibroblast activation (**Figure 4 D**), but PTEN knockdown on soft hydrogels resulted in increased levels of αSMA **(Figure 4 E,F)**. On stiff hydrogels, we observed lowered levels of PTEN with higher levels of αSMA (**Supplementary Figure 8 A-C**). However, the effect of PTEN on VICs cultured on stiff hydrogels was much lower compared to soft hydrogel conditions, as VICs are already activated on stiff hydrogels, but the effects observed were statistically significant **(Supplementary Figure 8 B,C)**. After confirming that PTEN protein levels were lower with our shRNA constructs, we next quantified mRNA expression levels of known fibrotic genes **(Figure 4 G)**. Compared to scrambled shRNA controls, PTEN knockdown VICs displayed higher levels of ACTA2, and COL1A1 expression. Collectively, these results further validate PTEN’s role as a regulator of the fibroblast-to-myofibroblast transition in VICs, using hydrogels as a mechano-regulator to introduce quiescence or activating matrix signals.

### High nuclear/cytoplasmic ratio of PTEN correlates with low αSMA levels in VIC on stiff hydrogels

In addition to the canonical role of PTEN in dephosphorylating PIP3 at the cell membrane, additional roles of PTEN and its nuclear localization have been recently identified, namely that PTEN is able to act as a transcription factor and alter global chromatin architecture and gene expression levels.^39^ Further, we have previously shown that matrix stiffness can influence the persistence of VIC myofibroblast activation through epigenetic remodeling, so we next sought to test whether nuclear PTEN is involved in this process. Specifically, we used quantitative image analysis to investigate the subcellular localization of PTEN in VICs cultured on both soft and stiff hydrogels.

Initially, we quantified αSMA levels of VIC cultures on soft and stiff hydrogels and separated the population into two classes: fibroblasts and myofibroblasts based on the intensity of αSMA signal. We observed that the quiescent VIC fibroblasts had a significantly higher nuclear to cytoplasmic PTEN ratio compared to myofibroblasts cultured on stiff hydrogels, where PTEN remained largely cytoplasmic **(Figure 5 A,B)**. We observed no statistically significant differences in the nuclear/cytoplasmic PTEN ratio between fibroblasts and myofibroblasts in VICs cultured on soft hydrogels, where VICs are quiescent. (**Figure 5 C-D**). Together, these results suggest that PTEN may serve an additional role in the nucleus, protecting against stiffness-induced matrix activation of myofibroblasts.

**Figure 5:**
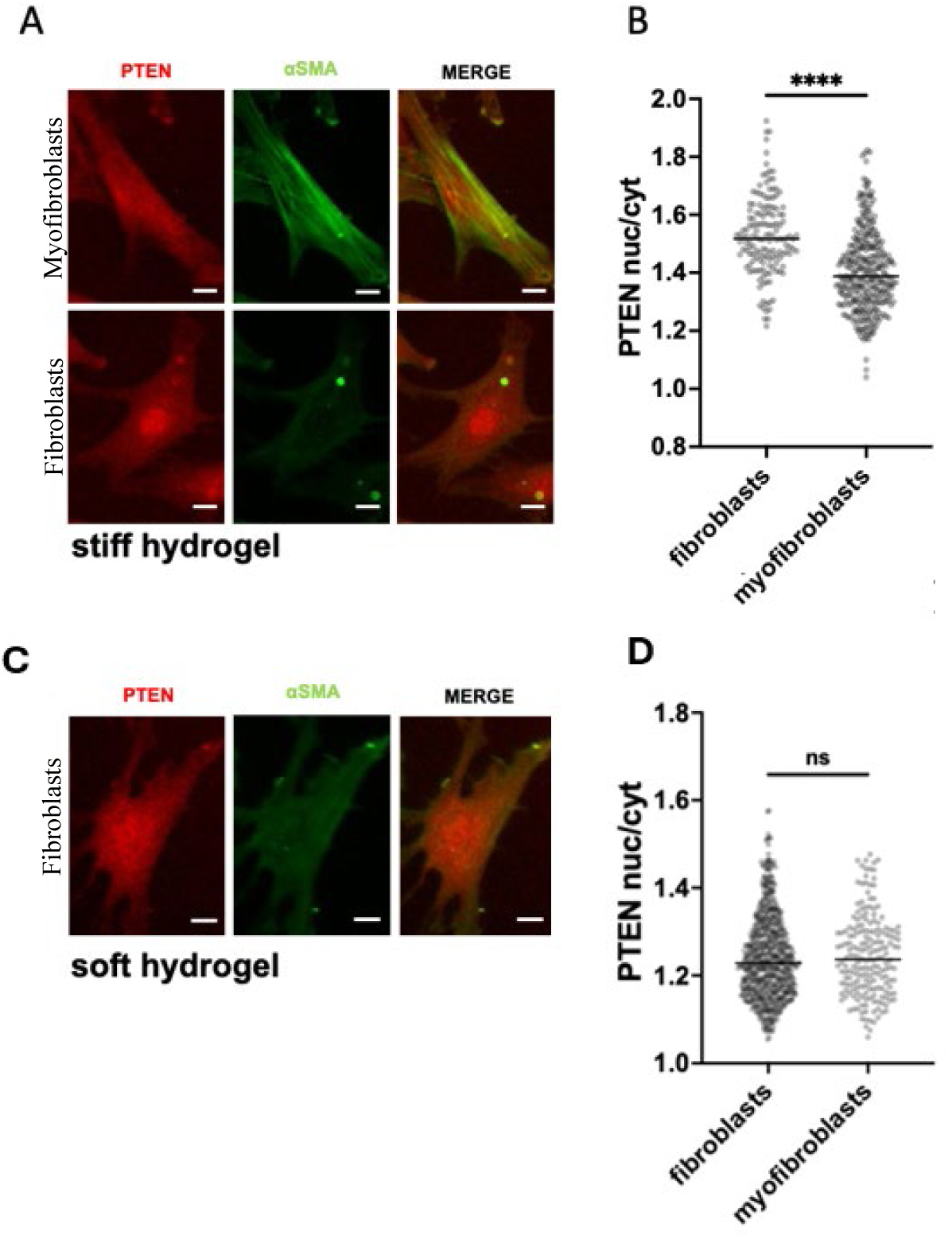
High nuclear/cytoplasmic ratio of PTEN correlates with low αSMA levels on stiff hydrogels. A. Single cell images of VICs cultured on stiff hydrogels for 3 days αSMA (green) and PTEN (red). Scale bar = 10 µm. B. Quantification of PTEN nuclear to cytoplasmic ratio in fibroblast and myofibroblast populations on stiff hydrogels (n>250 cells from 4 hydrogels, one-way ANOVA test, ****p<0.001). C. Single cell images of VICs cultured on soft hydrogels for 3 days αSMA (green) and PTEN (red). Scale bar = 10 µm. D. Quantification of PTEN nuclear to cytoplasmic ratio in fibroblast and myofibroblast populations on soft hydrogels (n>250 cells from 4 hydrogels, one-way ANOVA test, ****p<0.001). E. Quantification of nuclear area of cells with high and low nuclear to cytoplasmic PTEN ratio on soft hydrogels (n>250 cells from 4 hydrogels).

To verify the role of PTEN localization in myofibroblast activation, we ectopically expressed PTEN in VICs with two distinct nucleotide sequences that signal either nuclear (nuclear localization signal-NLS) or cytoplasmic (nuclear exclusion signal-NES) localization. Both plasmid constructs contain a hemagglutinin (HA) tag to verify transfection efficiency. VICs were plated on stiff 2D hydrogels and transfected with either NES PTEN or NLS PTEN plasmids with ∼4-7% efficiency **(Figure 6 D)**. After 3 days, we observed increased PTEN nuclear/cytoplasmic ratio in the NLS PTEN condition compared to NES PTEN and untransfected control, indicating that the transfected plasmid constructs localized PTEN as expected **(Figure 6 A, B)**. More importantly, αSMA was significantly downregulated among the cells transfected with NLS PTEN, while αSMA levels were slightly upregulated in the NES PTEN compared to untransfected control **(Figure 6 A, C)**, indicating that nuclear PTEN plays a protective role against myofibroblast activation on stiff hydrogels.

**Figure 6:**
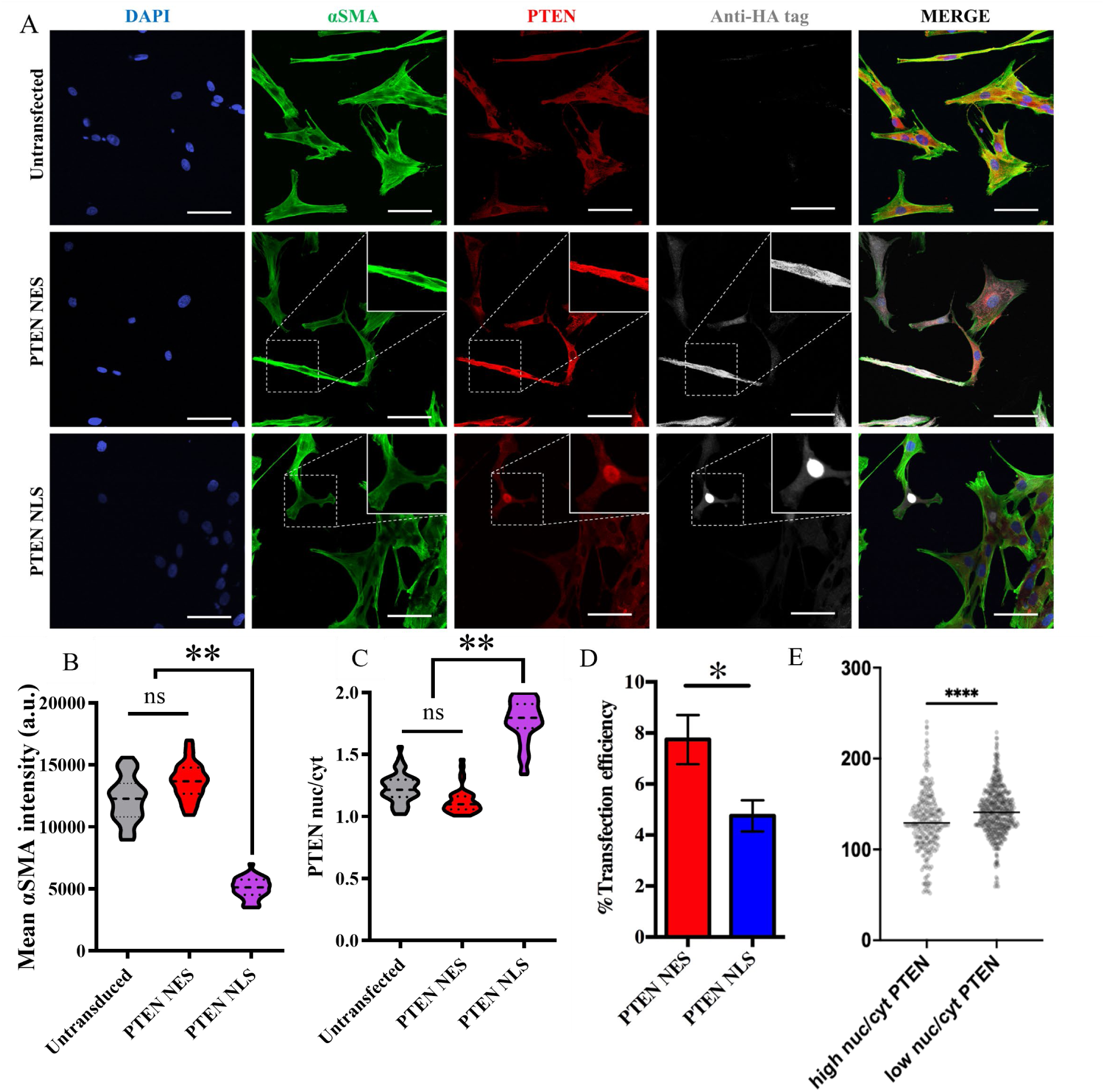
Ectopic expression of NLS PTEN downregulates *α*SMA on stiff hydrogels. **A.** Single cell images of VICs cultured on stiff hydrogels for 3 days αSMA (green), PTEN (red) and anti-HA tag (white). Scale bar = 10 µm. **B.** Quantification of αSMA among untransfected VICs and VICs with ectopic expression of NES and NLS PTEN on stiff hydrogels. (n=40 cells from 6 hydrogels, one-way ANOVA test, ****p<0.001) **C.** Quantification of the ratio of nuclear to cytoplasmic PTEN among untransfected, NES and NLS VICs (n=40 cells from 6 hydrogels, one-way ANOVA test, ****p<0.001). **D.** Quantification of the % transfection efficiency between NES and NLS PTEN. (n=40 cells from 6 hydrogels, one-way ANOVA test, ****p<0.001) **E.** Quantification of nuclear area of cells with high and low nuclear to cytoplasmic PTEN ratio on stiff hydrogels (n>250 cells from 4 hydrogels).

As nuclear PTEN has been shown to play a role in nuclear dynamics and epigenetic modifications, we wanted to test if there was a correlation between high nuclear PTEN and nuclear area in VICs.^39^ For VICs cultured on stiff hydrogels, we separated the cell population into two cohorts based on their nuclear to cytoplasmic PTEN ratio. VICs with a high nuclear/cytoplasmic PTEN ratio (nuclear/ cytoplasmic PTEN ≥ than 1.5) had significantly smaller nuclei compared to VICs with low nuclear/cytoplasmic PTEN ratio (nuclear/ cytoplasmic PTEN < than 1.5) **(Figure 6 E)**. This observation lead us to a more in depth investigation of the role of PTEN in chromatin dynamics.

Since the nuclear/cytoplasmic ratio of PTEN affected nuclear size in addition to αSMA levels, we subsequently studied the effect of the localization of PTEN on VIC epigenetic modifications. PTEN has been implicated in the regulation of histone activity in the context of fibrosis and heart failure, thus, we were interested in the correlation between the protective role of PTEN against αSMA upregulation and the levels of the tri-methylation of histone 3 on lysine-9 (H3K9me3) on VICs cultured for 3 days on soft and stiff hydrogels transfected with NES PTEN or NLS PTEN. Consistent with our previous results, the high nuclear/cytoplasmic ratio of PTEN, induced by NLS PTEN transfection, downregulated αSMA on stiff hydrogels but also upregulated H3K9me3, increasing its nuclear levels. In the NES PTEN as well as the untransfected control conditions, H3K9me3 levels remained significantly lower (**Figure 7A-D**). Interestingly, on soft hydrogels, where VICs display low αSMA levels, the increase in nuclear/cytoplasmic PTEN ratio, similarly, resulted in increased H3K9me3 levels **(Supplementary Figure 9A-D)**. This, indicated that the PTEN-H3K9me3 upregulation modulates αSMA levels, but their interactions are independent of VIC myofibroblast activation.

**Figure 7:**
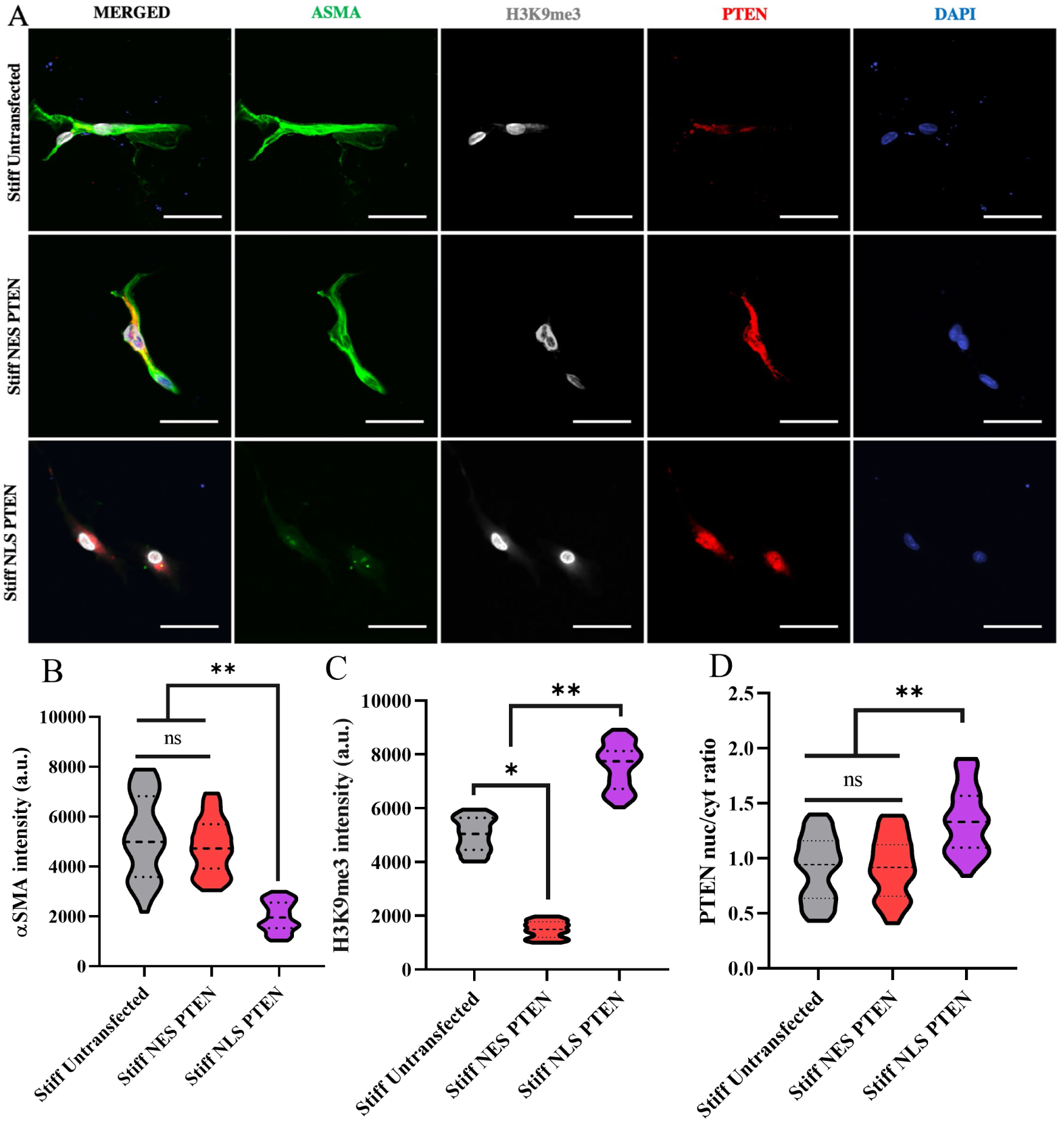
Nuclear localization of PTEN upregulates H3K9me3 on stiff hydrogels and protects against myofibroblast activation. **A.** Single cell immunocytochemical images of untransfected VICs and VICs with ectopic expression of NES and NLS PTEN cultured on stiff hydrogels for 3 days stained for αSMA (green), PTEN (red) and H3K9me3 (white). Scale bar = 10 µm. **B.** Quantification of αSMA intensity among untransfected VICs and VICs with ectopic expression of NES and NLS PTEN on stiff hydrogels. (n=50 cells from 4 hydrogels, one-way ANOVA test, ****p<0.001) **C.** Quantification of H3K9me3 nuclear intensity. (n=50 cells from 4 hydrogels, one-way ANOVA test, ****p<0.001) **D.** Quantification of the ratio of nuclear to cytoplasmic PTEN among untransfected, NES and NLS VICs (n=50 cells from 4 hydrogels, one-way ANOVA test, ****p<0.001).^40–42^

Finally, we were interested in the translation of our *in vitro* results of the PTEN-H3K9me3 interactions on human healthy and fibrotic heart valve tissue. Interestingly, histological analysis positively correlated the levels of PTEN with the levels of H3K9me3 in healthy tissue and inversely correlated with αSMA levels in fibrotic tissue (**Figure 8 A-C**). Colocalization analysis of H3K9me3 with DAPI, confirmed its nuclear localization albeit H3K9me3 only partially localizing with PTEN in human tissues **(Figure 8 D)**, suggesting an important but complex role of PTEN in the regulation of αSMA in human heart valve tissue.

**Figure 8:**
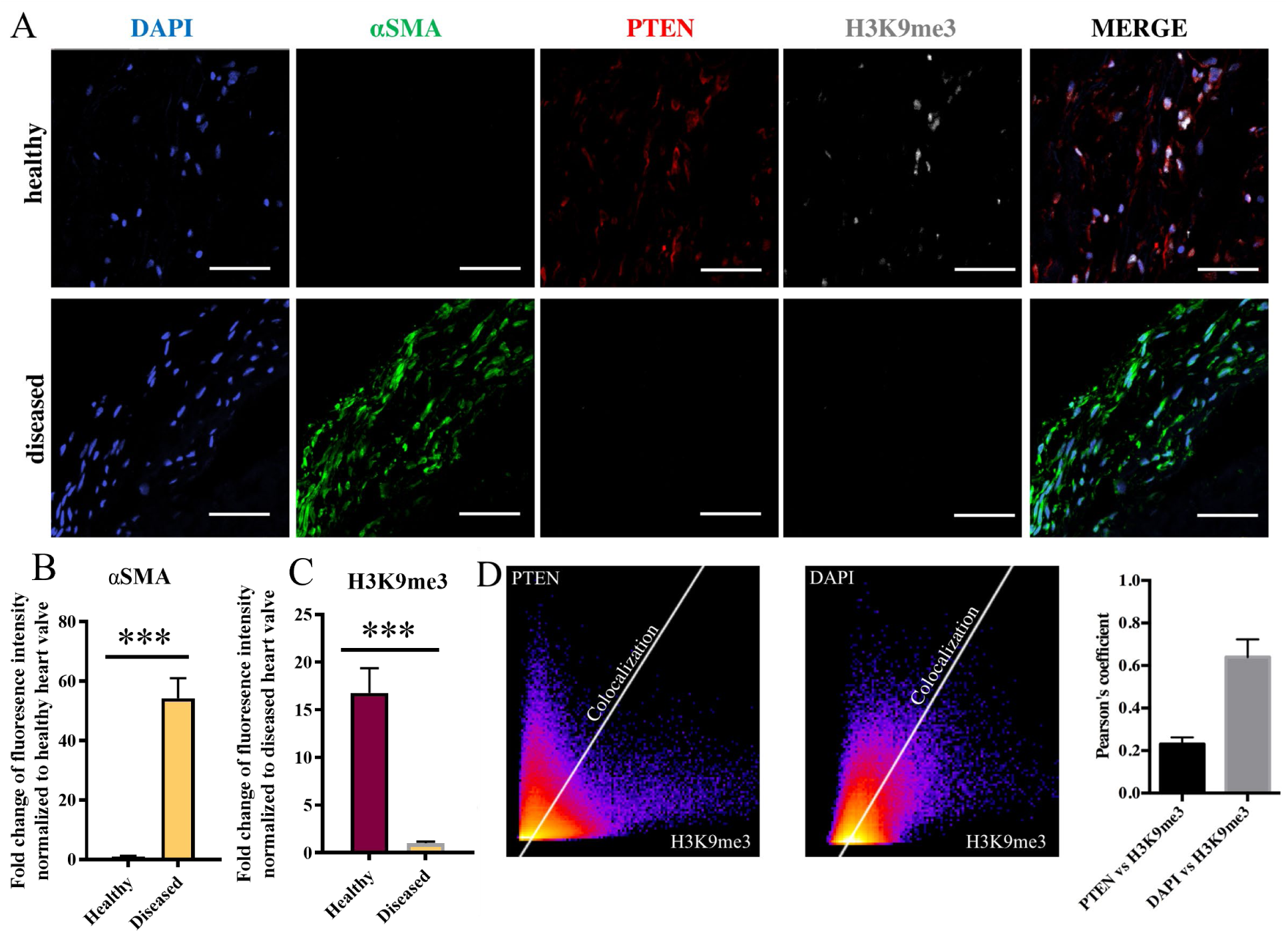
PTEN levels correlate with H3K8me3 levels in healthy human heart valve tissue while inversely correlating with *α*SMA on diseased tissue. **A.** Representative histological images of AVS and healthy human heart valve tissue (male) stained for DAPI (blue), αSMA (green), PTEN (red) and H3K9me3 (white) levels in valvular interstitial cells. Scale bar = 40 µm **B.** Quantification of average αSMA intensity (normalized to DAPI per mm^2^) for healthy and diseased tissue. (normalized to DAPI per mm^2^, n=5 fields of view per tissue, 2 biological replicates, t-test, ***p<0.001). **C.** Quantification of average H3K9me3 intensity (normalized to DAPI per mm^2^) for healthy and diseased tissue. (normalized to DAPI per mm^2^, n=5 fields of view per tissue, 2 biological replicates, t-test, ***p<0.001). **D.** Colocalization analysis of H3K9me3 and DAPI and αSMA. Plot of the classical Pearson’s coefficient. (n=5 fields of view per tissue, 2 biological replicates)

## Discussion

Results presented herein suggest that PTEN regulates myofibroblast activation in VICs, which is consistent with the emerging notion that PTEN plays a central protective role against fibrosis development in numerous tissues and organ systems.^25–27,29,36^ as well as myogenic differentiation of smooth vascular smooth muscle cells^62^. In line with this, we observed decreased levels of PTEN in diseased human valve leaflet tissues compared to healthy ones. The results linking PTEN with aortic valve stenosis complement what has been shown in tissue samples from patients with pulmonary fibrosis, where an inverse correlation of the expression of PTEN and αSMA was observed.^43^ Further, loss of PTEN protein and downregulation in its phosphatase activity have both been linked to development of pulmonary fibrosis.^44–48^ To further investigate the role of PTEN, we exploited biomaterial matrices to study the cross-talk between mechanosensing and matrix signaling, in the absence of other confounding factors, that can influence the VIC fibroblast-to-myofibroblast transition. We focused on elucidating how regulation of PTEN activity alters VIC activation/quiescence in both soft (healthy) and stiff (fibrotic-like) hydrogel microenvironments. Our findings suggest that PTEN has multiple modes of action depending on PTEN localization and matrix stiffness (**Figure 9**). We observe total PTEN expression levels decrease with matrix stiffness, inverse to the effects of stiffness on VIC myofibroblast activation. However, the spatial distribution of PTEN matters, as high nuclear to cytoplasmic ratio of PTEN on stiff hydrogels correlates with the quiescent fibroblast phenotype, as observed in **Figure 5** and may protect against myofibroblast activation when VICs are grown in stiff, fibrotic-like microenvironments.

**Figure 9:**
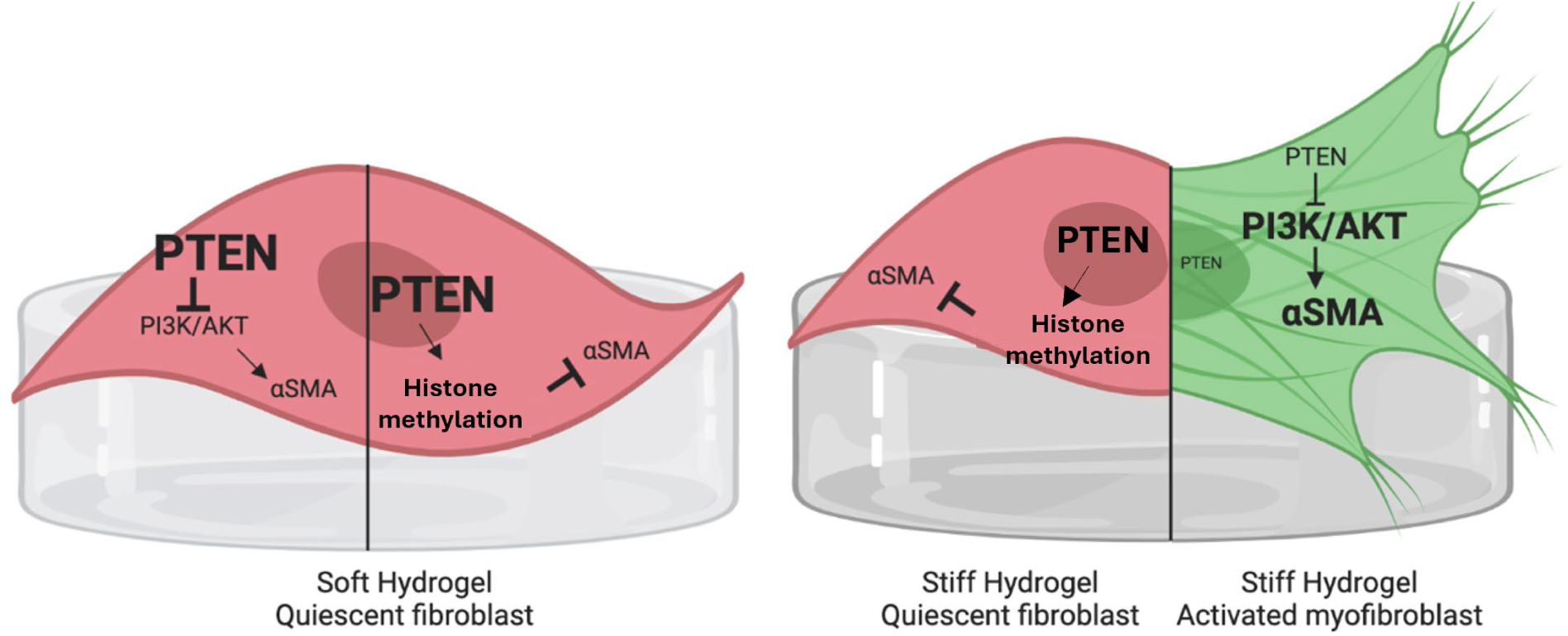
Schematic of PTEN activity in VICs on soft and stiff hydrogels. **A.** VICs cultured on soft hydrogels are quiescent due to the high PTEN activity in both the cellular cytosol and nucleus. Cytosolic PTEN acts as an antagonist of PI3K/AKT, while nuclear PTEN is involved in the regulation of histone expression. **B.** VICs differentiate towards myofibroblasts on stiff hydrogels due to the downregulation of PTEN, resulting in increased levels of phosphor-PI3K/AKT and elevated αSMA expression levels. However, when PTEN is ectopically expressed in the nucleus, αSMA levels are downregulated through increased expression of H3K9me3, maintaining VIC quiescence.

Further, we show that PTEN directly influences myofibroblast activation in VICs. PTEN is a known inhibitor of the PI3K/AKT pathway, and results from our lab reported by Wang *et al.* have shown that the phosphorylation of PI3K/AKT pathway is central to matrix stiffness induced myofibroblast activation in VICs.^24^ We validate the involvement of AKT in myofibroblast activation by pharmacological inhibition of phospho-AKT through MK-2206, which resulted in decreased αSMA levels. This led us to examine the relative role of PTEN, an AKT antagonist, abrogating the effects of mechanosensing. Namely, PTEN activators decrease, while PTEN inhibitors increase myofibroblast activation, respectively. Furthermore, in addition to being regulated by matrix stiffness, PTEN levels are also epigenetically regulated. We have demonstrated increased mRNA expression of PTEN by 5-azacytidine and itraconazole treatment (**Figure 3**, **Supplementary Figure 5**). Epigenetic silencing of PTEN has been proposed by methylation of the CpG islands within the promoter, which is involved in several cancers.^49–53^ As such, methyltransferase inhibitors 5-azacytidine and decitabine have been previously identified as PTEN activators by blocking methylation of the promoter and increasing gene expression.^28,54^ Interestingly, global methylation levels are reported to increase in AVS, which suggests that DNA methylation inhibitors might also be used to reverse this process.^55^

While relative cell PTEN levels are strong indicators of myofibroblast activation in this study and others, subcellular localization of PTEN has also been shown to play important roles in influencing cell phenotype.^43,56–58^ We observe that high nuclear localization of PTEN correlates with the quiescent fibroblast phenotype and smaller nuclei for VICs on stiff gels, but no correlations were observed with VICs on soft hydrogels. This suggests high nuclear localization of PTEN may protect against myofibroblast activation. Although loss of PTEN has been associated with fibrosis, subcellular localization of PTEN to the nucleus has not been previously studied in relation to fibrotic disease. To investigate the role of PTEN nuclear localization in myofibroblast activation, we ectopically expressed NES and NLS PTEN in VICs grown on stiff substrates, where nuclear PTEN significantly downregulated αSMA expression. A high level of nuclear PTEN, regulating epigenetic modifications, might suggest resistance to the transient myofibroblast state and continued protection against activation.

Complementing this observation, recent work from our lab linked histone regulation and chromatin condensation with myofibroblast activation.^19,59^ Specifically, Walker *et al.* used ATAC-seq to show that transient myofibroblasts have a more open chromatin structure, which subsequently condenses again to the stabilized phenotypic condition.^59,19^ PTEN has been shown to interact with chromatin by directly binding to histone H1 and controlling epigenetic modifications^60^, while nuclear PTEN levels correlate with H3K9me3 intensity, a marker for chromatin condensation, and depletion of H3K9me3 is observed in PTEN knockout cells.^56^ Further, our results are consistent with PTEN levels *in vivo* that correlate with H3K9me3 levels and potentially a protective effect against αSMA upregulation and fibrosis. Mechanistically, PTEN may reinforce the H3K9me3 suppression of the PPARγ gene, decreasing the expression of fibrotic markers such as αSMA and COL1.^61^ Taken together, we propose that nuclear PTEN promotes the VIC quiescent fibroblast phenotype by indirectly controlling αSMA expression at an epigenetic level. However, the mechanism that leads to the heterogeneity of nuclear PTEN observed in the VIC population remains a topic for future investigation.

In conclusion, we have shown that PTEN and its subcellular localization play an important role in regulating myofibroblast activation in VICs. Increased global levels of PTEN expression, either by low matrix stiffness or by pharmacological treatment, protect against myofibroblast activation. High levels of nuclear PTEN correlate with a H3K9me2 expression levels and the quiescent fibroblast phenotype, even on stiff hydrogel matrices. Importantly, this correlation is recapitulated in human healthy and fibrotic heart valve tissues, where we have observed mutually exclusive expression of αSMA and PTEN. Thus, we demonstrated that PTEN may be a valuable pharmacological target for non-invasive therapeutics for valve fibrosis and lead to further understanding of the dynamics of AVS progression.

## Acknowledgments

We thank Dr. Robert Weiss for valuable discussions and feedback. We acknowledge National Institutes of Health R01 HL142935 and R01 HL132353 awarded to KSA.

## Author contributions

D.B., G.T., and K.S.A. designed and performed all the experiments. B.K. and A.K. performed image analysis, mathematical analysis, and data 32 analysis. D.B., B.E.K., G.T., K.B., M.W.E. and K.S.A. contributed to writing and editing the paper.

## Supplementary Material

**Supplementary Figure 1:**
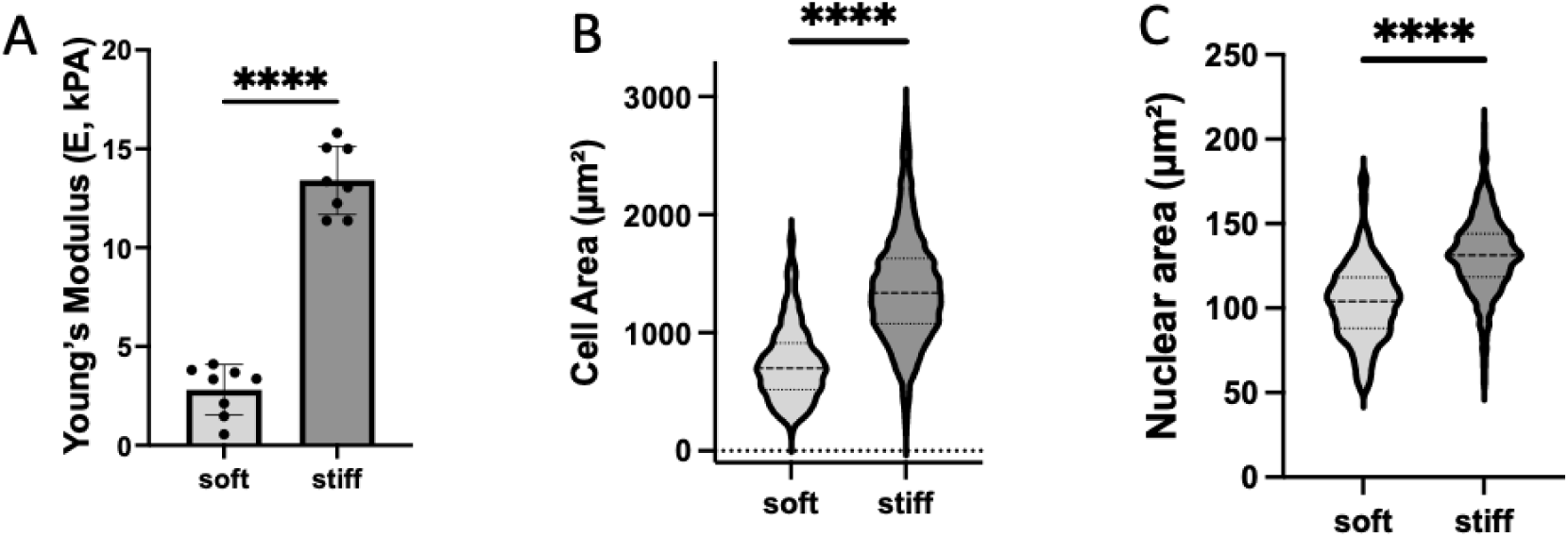
VICs display increased cellular and nuclear area on stiff hydrogels. **A.** Young’s modulus for synthesized soft and stiff swollen hydrogels by shear rheology. (n = 8, unpaired t test, ****p < .001). **B.** Quantification of cell size of VICs cultured on soft or stiff hydrogels for 3 days (n>740 cells from 4 hydrogels, unpaired t test, ****p<0.001). **C.** Quantification of nuclear area of VICs cultured on soft or stiff hydrogels for 3 days (n>740 cells from 4 hydrogels, unpaired t test, ****p<0.001).

**Supplementary Figure 2:**
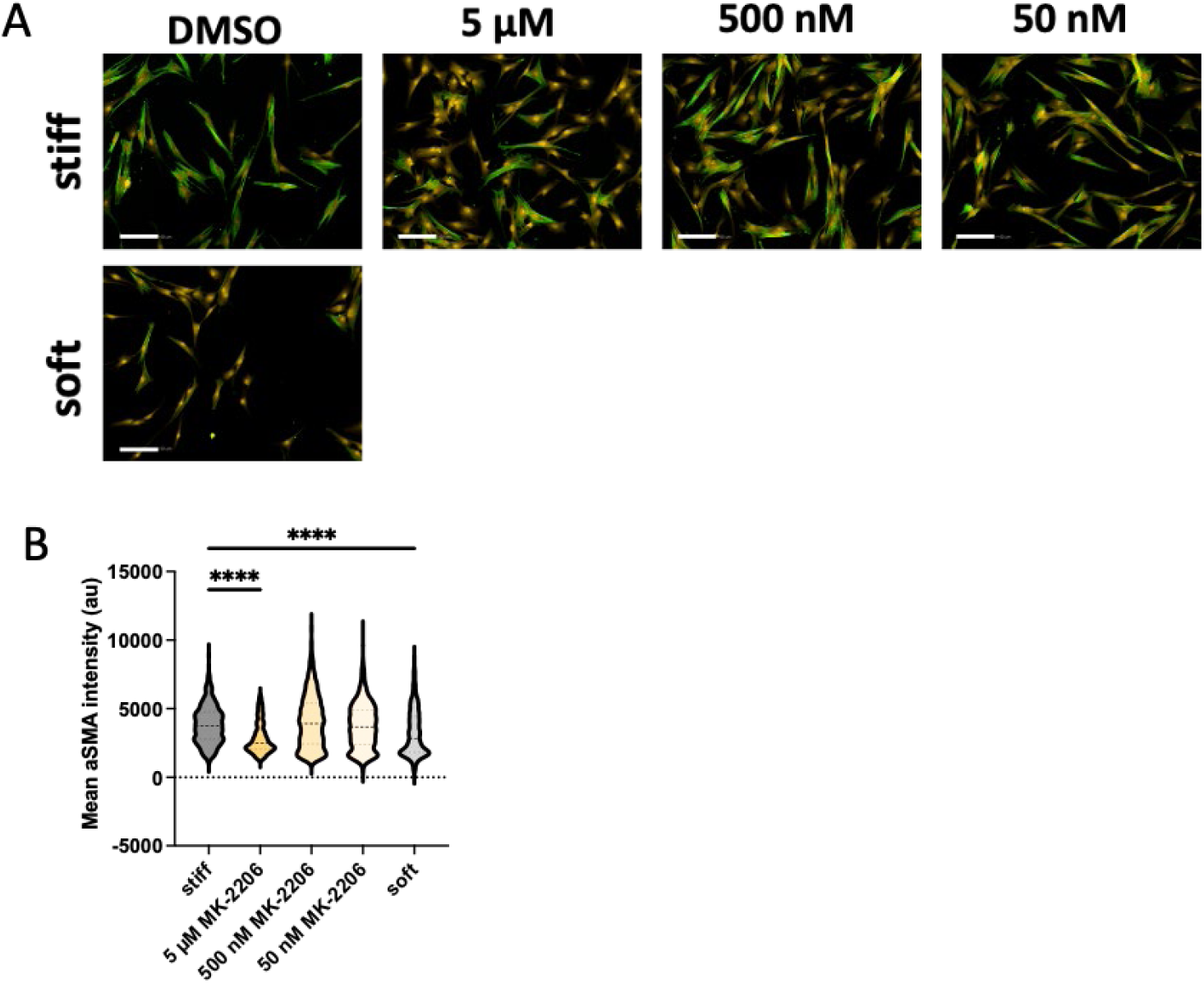
VICs display decreased myofibroblast activation with AKT inhibition. **A.** Representative images of VICs cultured on soft and stiff hydrogels for 3 days, αSMA (green) and cytoplasm (yellow). Scale bar = 100 µm **B.** Quantification of αSMA (green) intensity for A. (n>1000 cells from 4 hydrogels, one-way ANOVA test, ****p<0.001).

**Supplementary Figure 3:**
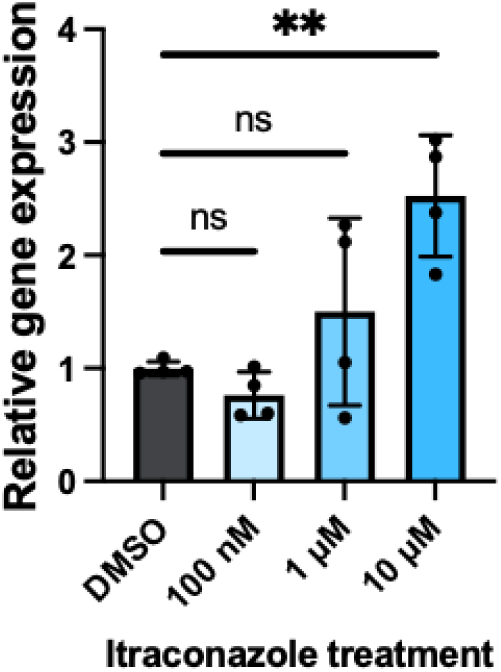
Itraconazole treatment increases PTEN gene expression. Relative mRNA expression levels of PTEN in VICs cultured on TCPS with treatment of vehicle or increasing concentration of PTEN activator itraconazole, n=4 wells, means ± SD shown, one-way ANOVA test, **p < .01.

**Supplementary Figure 4:**
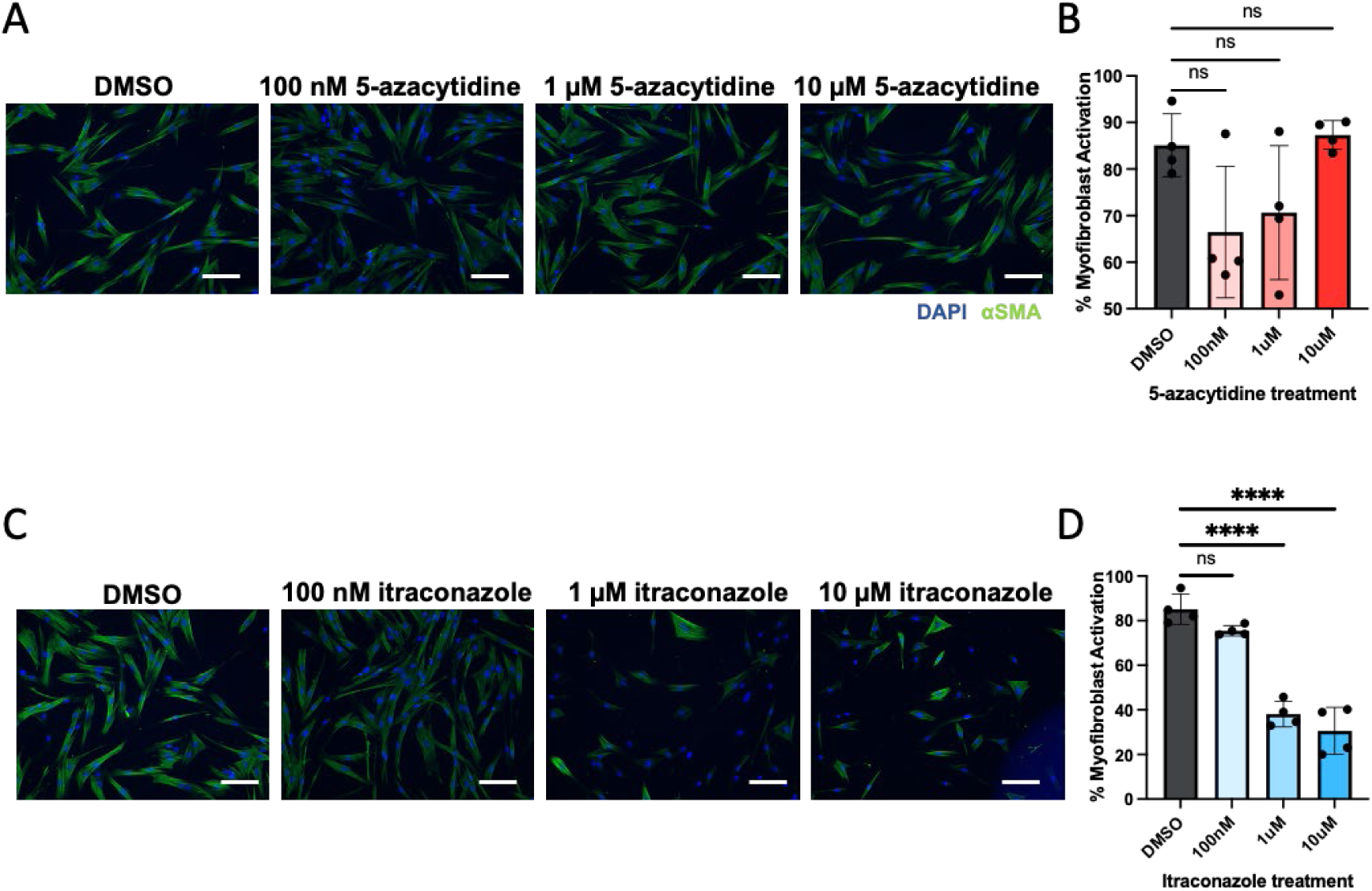
PTEN activators decrease myofibroblast activation in VICs. **A.** Representative images of VICs cultured on TCPS for 3 days with indicated concentrations of PTEN activators 5-azacytidine, αSMA (green) and nuclei(DAPI). Scale bar = 100 µm. **B.** Quantification of myofibroblast activation for A, n=4 wells, one-way ANOVA test. **C.** Representative images of VICs cultured on TCPS for 3 days with indicated concentrations of PTEN activators itraconazole, αSMA (green) and nuclei(DAPI). Scale bar = 100 µm. **D.** Quantification of myofibroblast activation for C, n=4 wells, one-way ANOVA test, ****p<0.001.

**Supplementary Figure 5:**
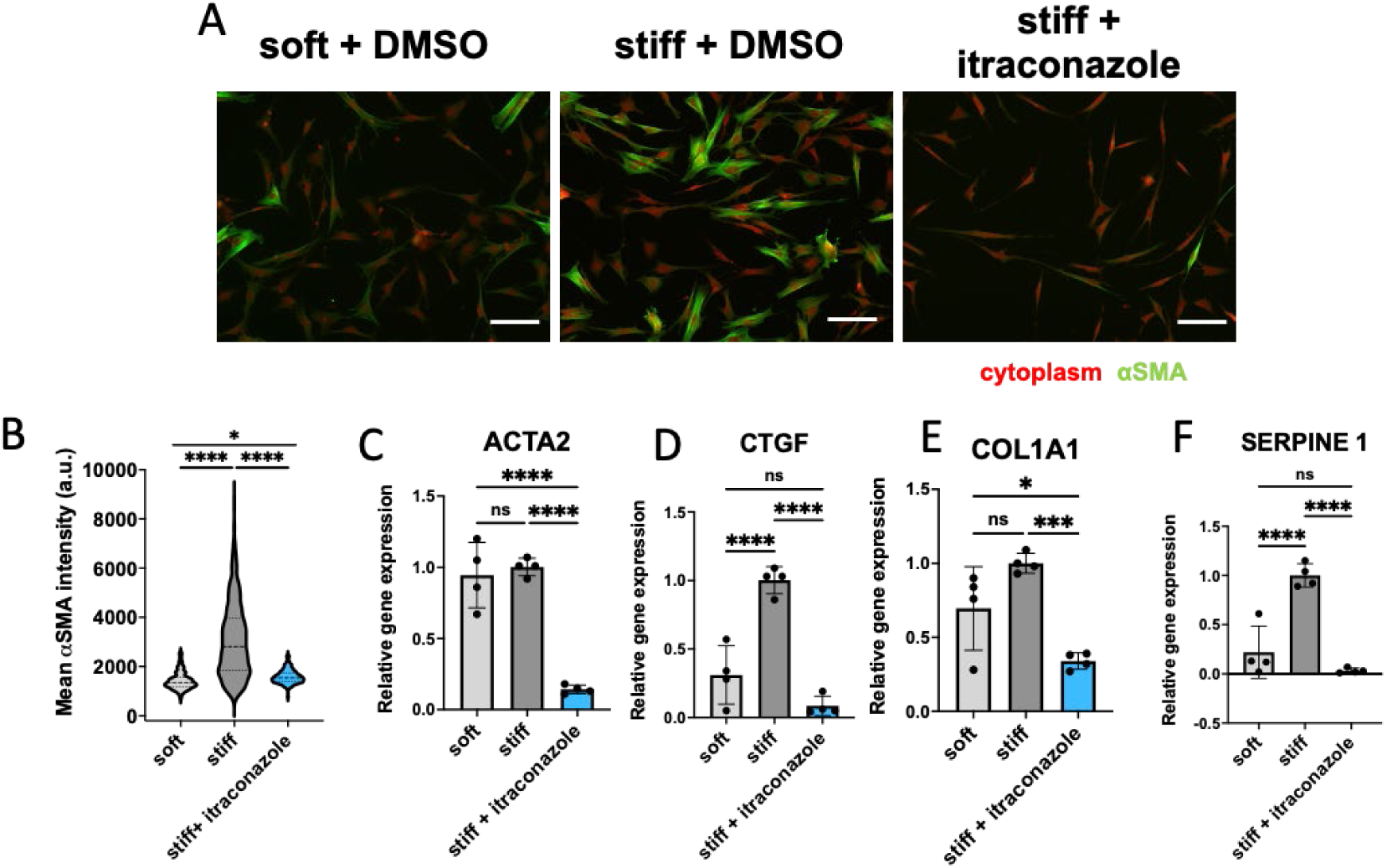
PTEN activators decrease myofibroblast activation in VICs. **A.** Representative images of VICs cultured on soft and stiff hydrogels for 3 days in the presence of DMSO vehicle or10 µM itraconazole (PTEN activator), αSMA (green) and cytoplasm(red). Scale bar = 100 µm. **B.** Quantification of αSMA intensity for A. (n>600 cells from 4 hydrogels, one-way ANOVA test, *p < .05, ****p<0.001). **C-F.** Relative mRNA expression levels of various fibrotic genes in VICs cultured on soft or stiff hydrogels treated with DMSO vehicle or PTEN activator itraconazole (10µM), n=4 hydrogels, one-way ANOVA test, *p < .05, **p < .01, ***p < .001,****p<0.001.

**Supplementary Figure 6:**
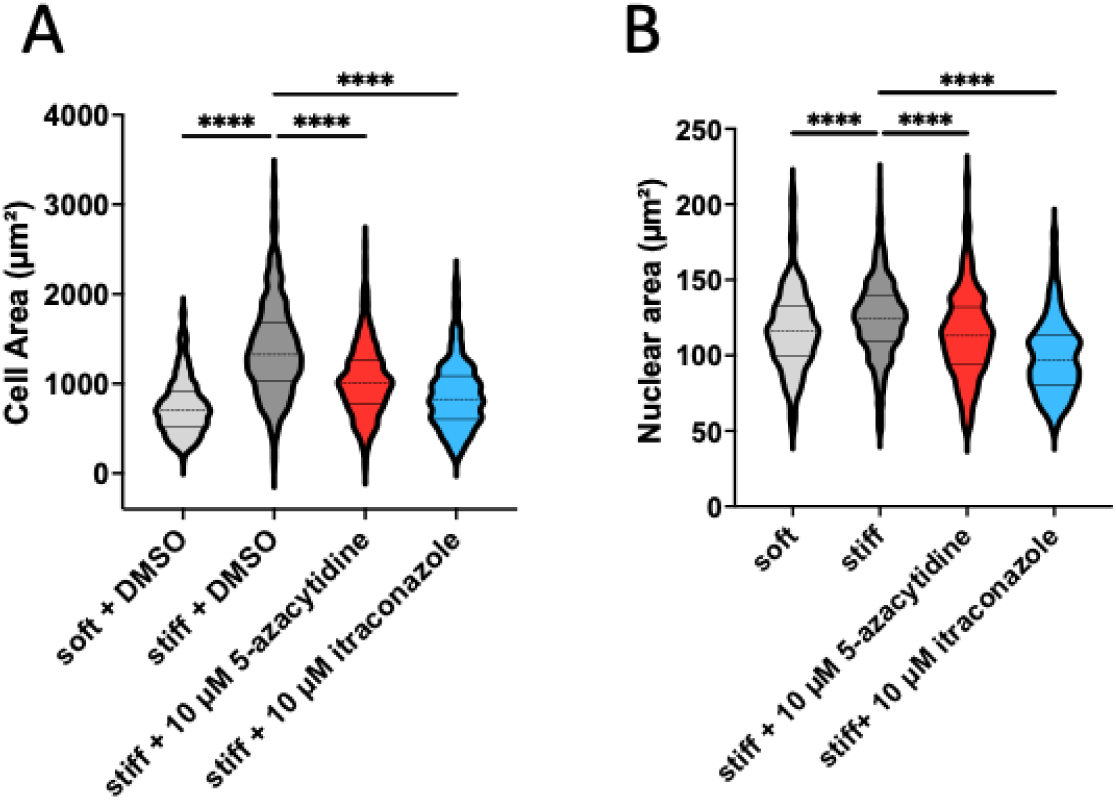
PTEN activators decrease cellular and nuclear area of VICs cultured on stiff hydrogels. **A.** Quantification of cell size of VICs cultured on soft or stiff hydrogels for 3 days (n>600 cells from 4 hydrogels, unpaired t test, ****p < 0.001). **B.** Quantification of cell size of VICs cultured on soft or stiff hydrogels for 3 days (n>600 cells from 4 hydrogels, unpaired t test, ****p < 0.001).

**Supplementary Figure 7:**
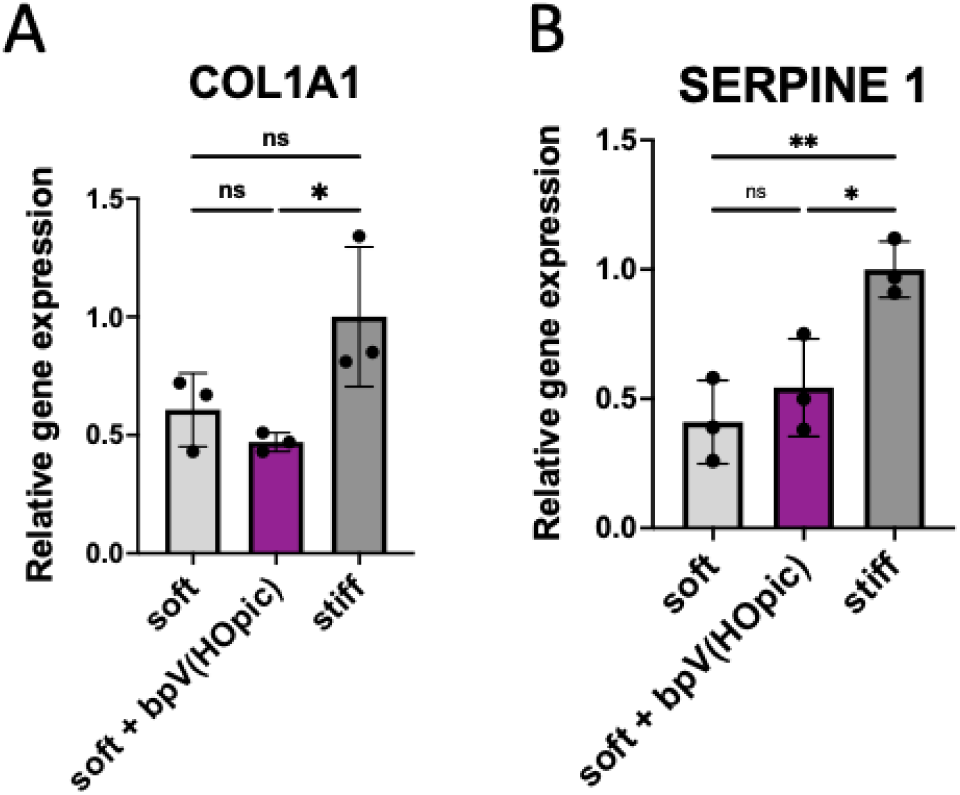
PTEN phosphatase inhibition does not increase mRNA levels of fibrotic genes A,B. Relative mRNA expression levels of various fibrotic genes in VICs cultured on soft or stiff hydrogels treated with DMSO vehicle or PTEN inhibitor bpV(Hopic) (100 nM), n=3 hydrogels, one-way ANOVA test, *p < .05, **p < .01.

**Supplementary Figure 8:**
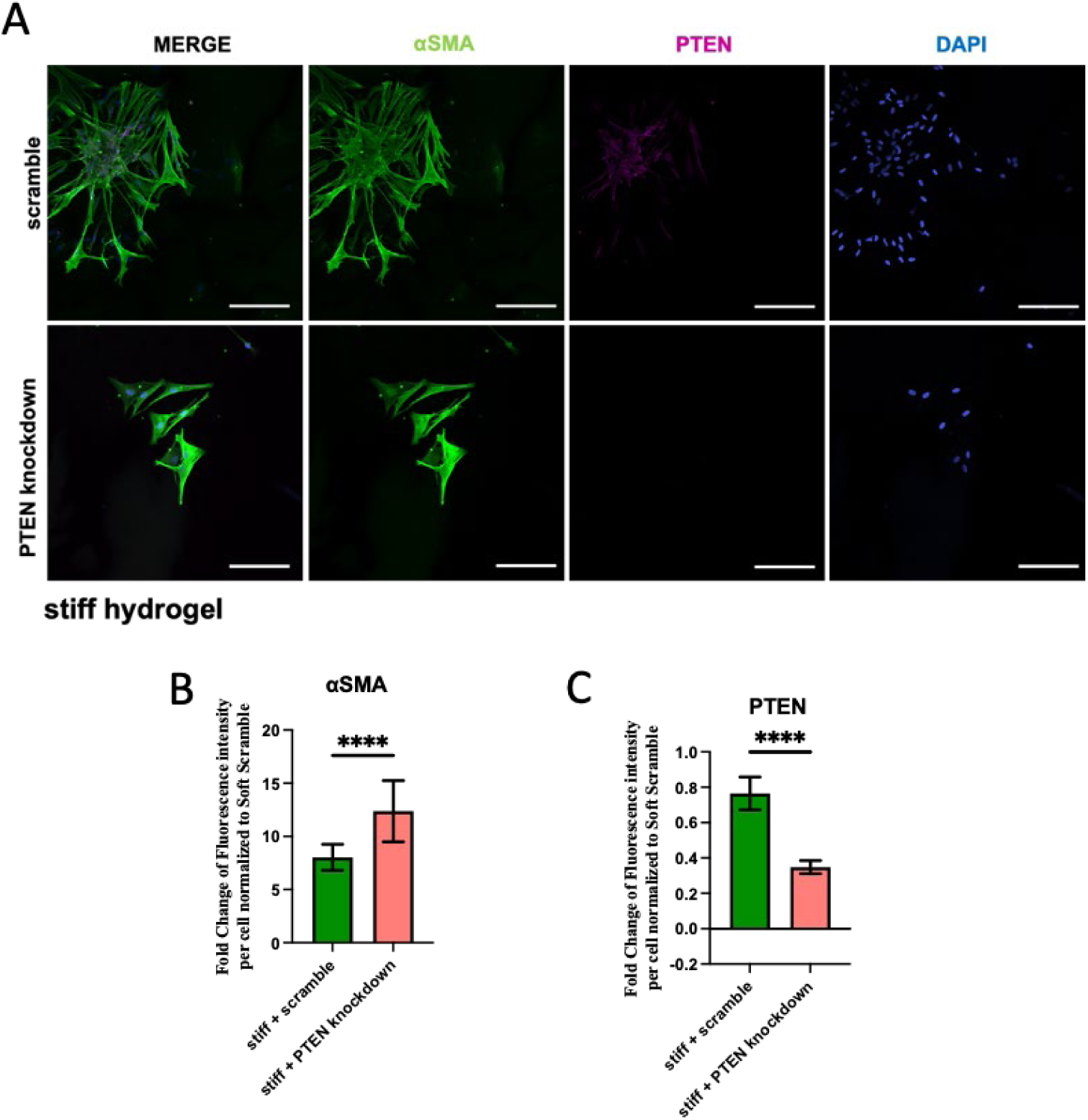
PTEN knockdown increases *α*SMA levels in VICs cultured on stiff hydrogels. **A.** Representative images of scrambled and PTEN knockdown VICs cultured on stiff hydrogels for 3 days, αSMA (green) and cytoplasm(red). Scale bar = 100 µm. **B.** Quantification of αSMA fluorescence intensity (n>30 cells, t-test, ****p<0.001) **C.** Quantification of PTEN fluorescence intensity (n>30 cells, t-test, ****p < 0.001).

**Supplementary Figure 9:**
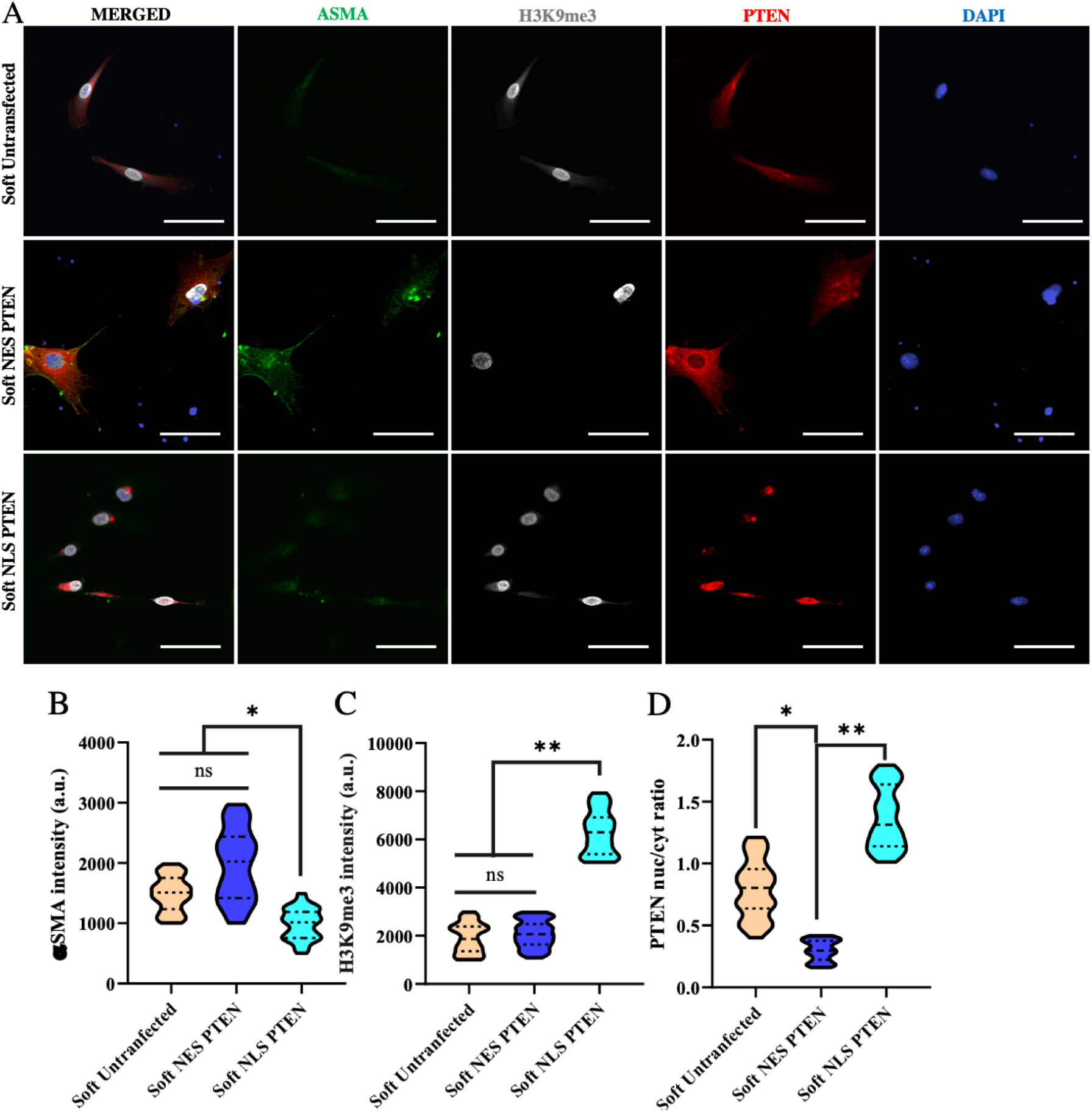
Nuclear localization of PTEN upregulates H3K9me3 on soft hydrogels. **A.** Single cell immunocytochemical images of VICs cultured on soft hydrogels for 3 days αSMA (green), PTEN (red) and H3K9me3 (white). Scale bar = 10 µm. **B.** Quantification of αSMA intensity among untransfected VICs and VICs with ectopic expression of NES and NLS PTEN on soft hydrogels. (n=50 cells from 4 hydrogels, one-way ANOVA test, ****p<0.001) **C.** Quantification of H3K9me3 nuclear intensity. (n=50 cells from 4 hydrogels, one-way ANOVA test, ****p<0.001) **D.** Quantification of the ratio of nuclear to cytoplasmic PTEN among untransfected, NES and NLS VICs (n=50 cells from 4 hydrogels, one-way ANOVA test, ****p<0.001).

**Table S1:**
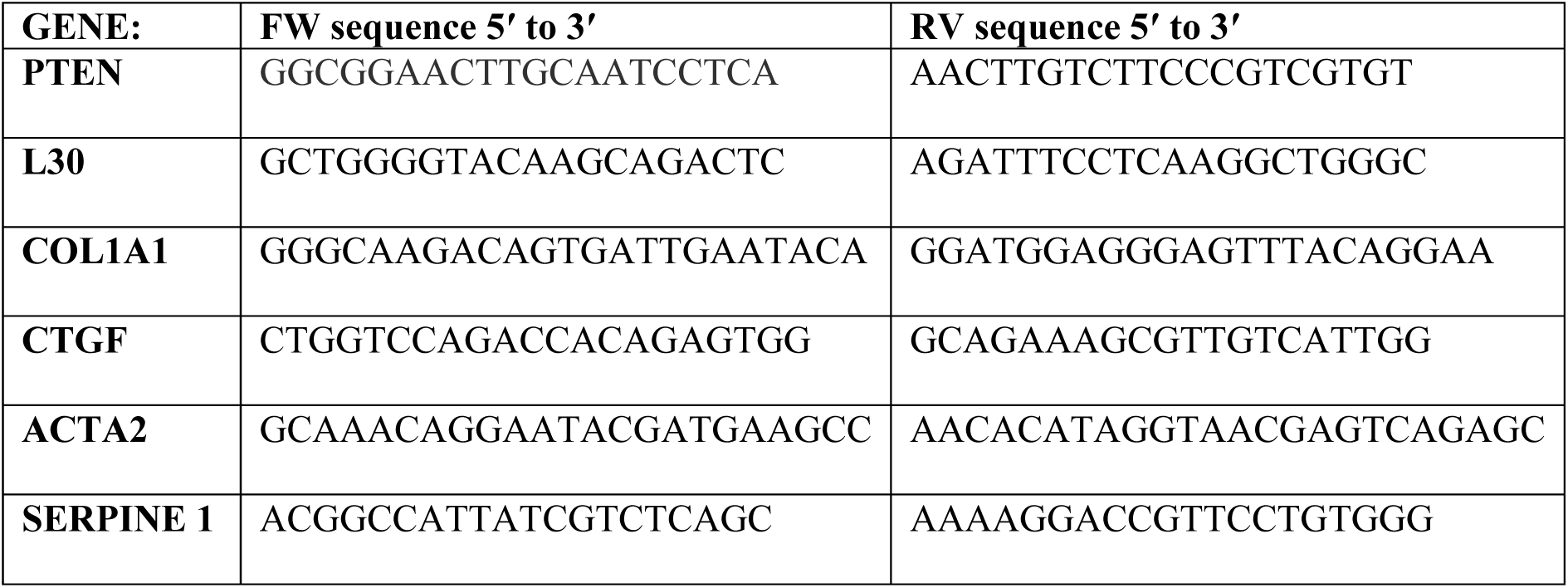
Custom Primers used for RT-qPCR.

**Table S2:** Patient Information of the tissue used.

| **Condition** | **Age** | **Sex** | **Race (Ethnicity)** | **Diagnosis** | **LVEF (%)** |
| --- | --- | --- | --- | --- | --- |
| **Healthy** | **45** | **M** | **american indian** | **non-ischemic** | **35** |
| **Stenotic** | **67** | **M** | **W** | **ischemic** | **29.5** |

